# Identification of viruses from fungal transcriptomes

**DOI:** 10.1101/2020.02.26.966903

**Authors:** Yeonhwa Jo, Hoseong Choi, Hyosub Chu, Won Kyong Cho

## Abstract

Viruses infecting fungi are referred to as mycoviruses. Here, we carried out *in silico* mycovirome studies using public fungal transcriptomes. We identified 468 virus-associated contigs assigned to five orders, 21 families, 26 genera, and 88 species. We assembled 120 viral genomes with diverse RNA and DNA genomes. The phylogenetic tree and genome organization unveiled the possible host origin of mycovirus species and diversity of their genome structures. Most identified mycoviruses were originated from fungi; however, some mycoviruses had strong phylogenetic relationships with those from insects and plants. The viral abundance and mutation frequency of mycoviruses were very low; however, the compositions and populations of mycoviruses were very complex. Although coinfection of diverse mycoviruses in the fungi was common in our study, most mycoviromes had a dominant virus species. The compositions and populations of mycoviruses were more complex than we expected. *Monilinia* viromes revealed that there were strong deviations in the composition of viruses and viral abundance among samples. *Gigaspora* viromes showed that the chemical strigolactone might promote virus replication and mutations, while symbiosis with endobacteria might suppress virus replication and mutations. This study revealed the diversity and host distribution of mycoviruses.

## INTRODUCTION

Viruses are small infective agents composed of nucleic acids including DNA or RNA that replicate in the infected host (1). Viruses are ubiquitous, presenting not only in most living organisms such as animals, plants, fungi, and bacteria but also in diverse environments (2–6).

Viruses do not have a conserved genetic element like bacteria and fungi (7). Moreover, isolation of pure viral particles is very difficult and challenging for the viruses with low titers and coinfecting viruses (8). The rapid development of next-generation sequencing (NGS) followed by bioinformatics tools has facilitated the identification of numerous eukaryotic microorganisms in diverse hosts and environments (9). In particular, the metatranscriptomics based on RNA-sequencing (RNA-seq) overcomes several obstacles associated with virus identification (10,11). NGS techniques enable us to identify both known and novel viruses in the host and provide several types of information, such as viral genomes and mutations (12,13).

Recently, virus-associated studies using NGS have used a new term, “virome,” which refers to a whole collection of viral nucleic acids in a specific host or a particular ecosystem (14,15). In fact, virome studies provide several types of information associated with viral communities, such as viral genomes, viral populations, mutations, phylogeny, evolution, recombination, and interaction of viruses with the host or ecosystem, which have been widely used in many such studies.

Viruses infecting fungi are referred to as mycoviruses (16). Most observed mycoviruses consist of double-stranded (ds) RNA genomes; however, observed mycoviruses with positive or negative single-stranded (ss) RNA and ssDNA genomes is gradually increasing (17,18). As compared to other viruses infecting animals and plants, most known mycoviruses lack an extracellular route for infection and movement proteins (19). Some of identified mycoviruses infecting phytopathogenic fungi have the ability to reduce the fungal hosts’ pathogenicity, known as hypovirulence (16). Many novel mycoviruses are now being identified based on metatranscriptomics, such as 66 novel mycoviruses with 15 distinct lineages from five plant-pathogenic fungi (20) and 59 viruses from 44 different fungi (21). Moreover, several studies have shown that fungal hosts are very often coinfected by various mycoviruses (21,22).

The rapid development and application of NGS techniques has increased the vast amount of sequencing data in public databases, such as the Short Read Archive (SRA) at NCBI, which are freely available. Many studies have demonstrated the usefulness of transcriptomic data for virus identification and virome study without any *a priori* knowledge of virus infection (21,23). Here, we carried out *in silico* bioinformatics analyses to identify viruses infecting fungi using available fungal transcriptome data and addressed several mycovirus-associated findings.

## MATERIALS AND METHODS

### Download of fungal transcriptomes

In order to identify virus-associated contigs (transcripts) in fungal transcriptomes, all available fungal assembled contigs were downloaded from the TSA database (https://www.ncbi.nlm.nih.gov/genbank/tsa/) using “fungi” as a query.

### Construction of viral genome and protein database

All viral genomes and protein sequences were downloaded from NCBI’s viral genome website (https://www.ncbi.nlm.nih.gov/genome/viruses/) on October 10, 2018. The downloaded sequences were used to set up a database for BLAST and DIAMOND using the following commands: “makeblastdb –in viral_genome.fasta –dbtype nucl –out viral_genome” and “diamond makedb –in viral_protein.faa -d viral_protein” (24).

### Identification of virus-associated contigs

All fungal transcripts were subjected to BLASTX search against the viral protein database using the DIAMOND program with E-value 1e-10 as a cutoff (diamond blastx -d viral_protein -q fungal_contigs.fasta -e 1e-10 -k 1 -f 6 qseqid sseqid pident length mismatch gapopen qstart qend sstart send evalue bitscore qlen -o fungal_contigs_viral_protein.txt). Out of 3,733,874 fungal transcripts, 7,244 contigs showing similarity to viral proteins were extracted for further analysis. The 7,244 extracted contigs were subjected to BLASTX using DIAMOND with E-value 1e-10 as a cutoff against NCBI’s NR database (ftp://ftp.ncbi.nlm.nih.gov/blast/db/FASTA/nr.gz) to remove sequences derived from hosts and other organisms such as eukaryotes and bacteria. We ultimately obtained 468 virus-associated contigs.

### Viral genome annotation

The 468 virus-associated contigs were again subjected to TBLASTX and BLASTN searches against the viral genome database to annotate the virus-associated contigs using E-value 1e-10. We selected virus-associated contigs larger than 1,000 bp for the prediction of open reading frames (ORFs) by the stand-alone version of ORFfinder in NCBI (ftp://ftp.ncbi.nlm.nih.gov/genomes/TOOLS/ORFfinder/linux-i64/). The predicted ORFs were compared to the related reference virus genome by BLASTP. Conserved domains in the virus-associated contigs were predicted by the SMART program (http://smart.embl-heidelberg.de/) (25).

### Phylogenetic tree construction

We obtained 153 viral protein sequences composed of 107 viral replicase-associated sequences, 17 CPs, 15 hypothetical proteins, six MPs, and 14 other proteins. We only used replicase-associated protein sequences as representatives of identified viruses for the phylogenetic analyses. To examine the phylogenetic relationship of identified viruses with known viruses, 107 viral replicase proteins were divided into 16 viral families, one viral order, and one unclassified group, resulting in 18 datasets. The viral replicase proteins in each dataset were subjected to BLASTP search against the NR database in NCBI. We included all matched protein sequences and removed redundant protein sequences.

Viral replicase sequences in each dataset were aligned using the L-INS-I method implemented in MAFFT version 7.450 (26). We removed all ambiguously aligned sequences by TrimAl (version 1.2) with the option “automated1” (27). We determined the best-fit model of amino acid substitution in aligned sequences using IQ-TREE (28). The maximum likelihood phylogenetic tree in each dataset was inferred using IQ-TREE with the ultrafast bootstrap method (29) and the SH-aLRT branch test (30). The obtained phylogenetic trees were visualized by FigTree (http://tree.bio.ed.ac.uk/software/figtree/).

### Estimation of viral abundance in fungal transcriptomes

We downloaded all raw sequence data associated with the 43 fungal transcriptomes containing virus-associated contigs from the SRA database in NCBI (https://www.ncbi.nlm.nih.gov/sra). Unfortunately, some raw sequences were not available. In many cases, there were several SRA datasets for a fungal transcriptome. We calculated viral abundance for an individual SRA dataset, not a single fungal transcriptome. The downloaded SRA data were converted into fastq files using the fastq-dump tool implemented in the SRA toolkit program (31). Each fastq file was mapped on the 468 virus-associated sequences using the Burrows-Wheeler Alignment (BWA) tool with the “BWA-MEM” algorithm (32). The mapped SAM file was subjected to pileup.sh in the BBMap program (https://sourceforge.net/projects/bbmap/) to extract the number of reads and FPKM values, which is a normalized estimation of gene expression considering the gene length and the sequencing depth in RNA-seq data. To calculate viral abundance in each fungal transcriptome, the number of virus-associated reads was divided by the total number of reads. We expected more reads from the large virus genomes and datasets with high sequencing depth. Therefore, we used FPKM values to calculate the proportion of viral abundance in a specific fungal transcriptome.

### Fungal transcriptome assembly and calculation of viral mutations

For the virome study, we selected two previously studied fungal transcriptomes, *Monilinia* transcriptomes including 18 libraries and *Gigaspora margarita* transcriptomes including 21 libraries. Both fungal transcriptomes were obtained from diverse fungal samples that were infected by many mycoviruses. However, the available assembled fungal transcriptomes were obtained by combining all libraries. Therefore, it was necessary to assemble an individual library for the virome study. Raw sequence reads from each condition were *de novo* assembled using the Trinity program (version 2.84) with default parameters (33). Then, each assembled transcriptome was subjected to BLASTN search against 468 virus-associated contigs. From each transcriptome, we extracted virus-associated contigs followed by BLASTX search against the NR database to remove non-viral sequences. Finally, we obtained virus-associated contigs from the individual condition. We mapped the raw sequence reads against the virus-associated contigs in each condition to calculate viral abundance and SNPs. Viral abundance in each condition was calculated by the number of viral reads and FPKM values.

### Identification of virus mutations

It is important to select proper reference sequences for the identification of virus mutations. For example, if we use a known reference viral genome for mapping, we can find several SNPs between two different viral genomes. However, the obtained SNPs are not viral mutations. For that, we used viral sequences as references and raw sequence reads in the identical condition for the calculation of SNPs. We obtained SNP results for 18 *Monilinia* and 21 *Gigaspora margarita* transcriptomes. SNPs were calculated as follows. For each condition, raw sequence reads were mapped on the viral sequences obtained from the identical condition using the BWA-MEM algorithm with default parameters. We converted the aligned reads in the SAM file format into the BAM file format using SAMtools (ver. 1.3) (34). After sorting and indexing the BAM file, SNP calling was performed using BCFtools (ver. 1.3) (35). We obtained a VCF (Variant Call Format) file containing information on SNPs representing virus mutations for each condition.

## RESULTS

### Identification of virus-associated contigs from fungal transcriptomes

To identify viruses from fungal transcriptomes, we obtained 3,733,874 fungal assembled contigs (4,268,333,264 bp) from 126 different fungal transcriptomes in the Transcriptome Shotgun Assembly (TSA) database (Fig. 1a and Table S1). Initially, all fungal contigs were subjected to a BLASTX search against the viral protein database, resulting in 7,244 contigs based on E-value 1e-10 as a cutoff. After removing sequences from hosts and other organisms, we obtained 468 virus-associated contigs (Fig. 1b and Table S2).

**Figure 1.**
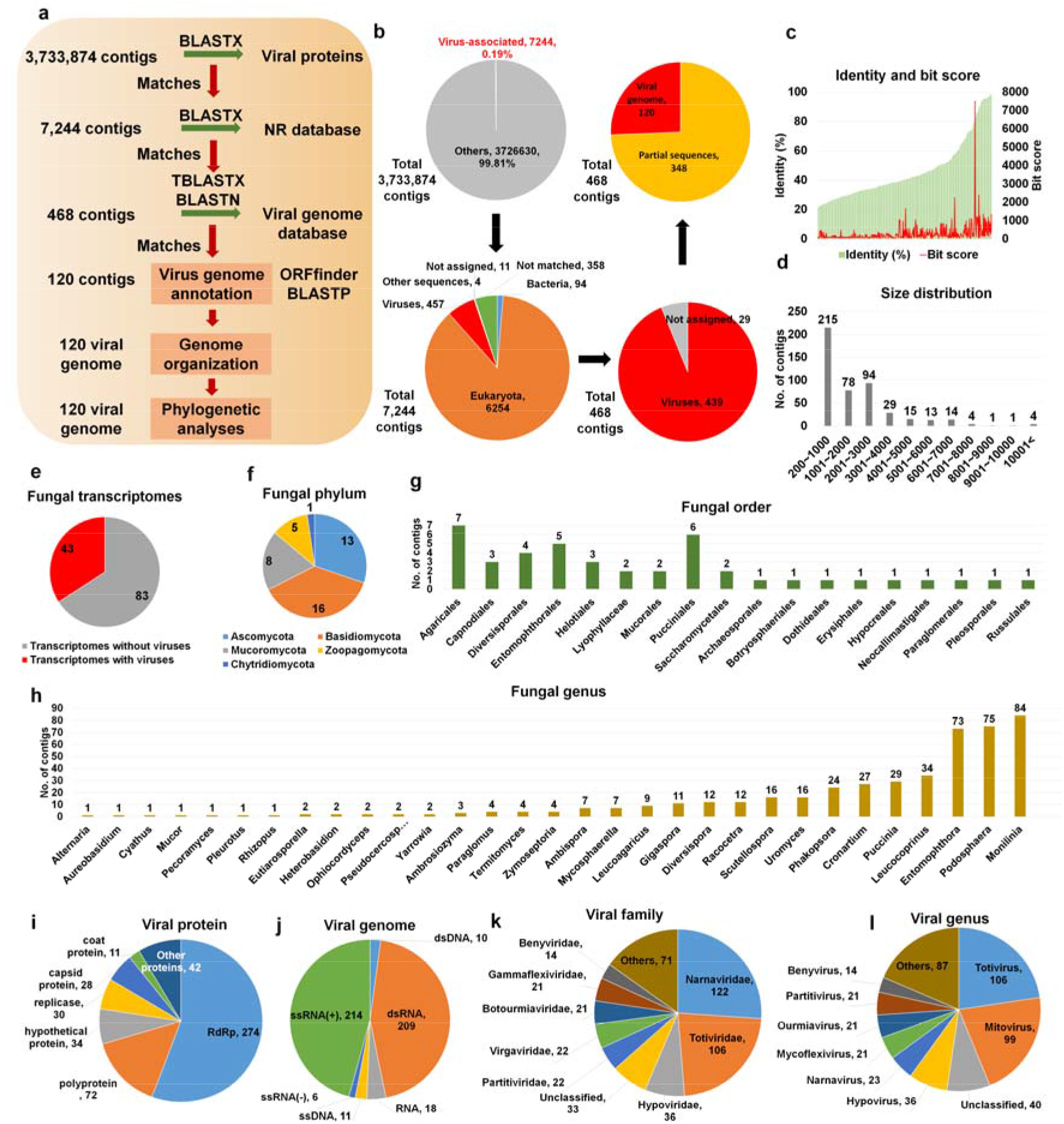
Identification of virus-associated contigs from fungal transcriptomes. (a) Schematic representation of bioinformatics workflow to identify virus-associated contigs from fungal transcriptomes. (b) Proportion of identified virus-associated contigs in each major step. (c) Distribution of protein identity (left) and bit score (right) for 468 virus-associated contigs. (d) Size distribution of 468 identified virus-associated contigs. (e) Number of fungal transcriptomes with and without viruses, respectively. Number of fungal transcriptomes containing viruses according to fungal phylum (f), order (g), and genus (h). Classification of virus-associated contigs according to viral proteins (i) and virus genome type (j). Classification of 468 virus-associated contigs according to viral family (k) and genus (l).

The BLAST results showed that the sequence identity (20.9–98.7%) and bit score (36.6–7520.6) for the 468 virus-associated contigs against viral reference genomes were relatively correlated (Fig. 1c). Most virus-associated contigs ranged from 201 to 1,000 bp (227 contigs) followed by 2,001 to 3,000 bp (97 contigs) (Fig. 1d). Moreover, five contigs were larger than 10 kb. Of the 126 fungal transcriptomes, only 43 included virus-associated contigs (Fig. 1e and Table S1). The 43 fungal transcriptomes were assigned to five fungal phyla, 18 orders, and 31 genera (Figs. 1f–1h). The most frequently identified fungal phyla were *Basidiomycota* (16 transcriptomes), *Ascomycota* (13 transcriptomes), and *Mucoromycota* (eight transcriptomes) (Fig. 1f). The top three fungal orders possessing viruses were *Agaricales*, *Pucciniales*, and *Entomophthorales* (Fig. 1g). At the genus level, *Monilinia*, *Podosphaera*, and *Entomophthora* were frequently infected by different viruses (Fig. 1h).

A majority of virus-associated contigs were homologous to known viral proteins, including RNA-dependent RNA polymerase (RdRp) (274 contigs) followed by polyprotein (72 contigs) and hypothetical protein (34 contigs) (Fig. 1i and Table S2). Most virus-associated contigs were derived from positive ssRNA genomes (214 contigs) and dsRNA genomes (209 contigs) (Fig. 1j). Furthermore, some contigs were derived from viruses with negative ssRNA, ssDNA, and dsDNA genomes.

The 468 virus-associated contigs were assigned to five viral orders, 21 families, 26 genera, and 88 species (Table S2). The frequently identified viral orders, families, and genera were *Tymovirales* (23 contigs) and *Picornavirales* (seven contigs); *Narnaviridae* (98 contigs), *Totiviridae* (95 contigs), and *Hypoviridae* (35 contigs) (Fig. 1k); and *Mitovirus* (84 contigs), *Hypovirus* (35 contigs), and *Totivirus* (16 contigs) (Fig. 1l), respectively.

### Viral genome assembly and phylogeny of novel mycoviruses

We assembled 120 viral genomes divided into 16 viral families, one order, and unclassified viruses based on obtained virus-associated contigs (Table S3). Most viruses were novel viruses with low sequence identity with known viruses and had a single genome, except for four viruses with two RNA segments. Except for one ssDNA virus, all viruses had RNA genomes, which were further divided into unknown RNA (12 viruses), dsRNA (46 viruses), positive ssRNA (57 viruses), and negative ssRNA genomes (four viruses). Of the 18 identified virus genera, many viruses (28) were unclassified, while totiviruses (20) and ourmiaviruses (13) were two major viral genera.

We analyzed the genome organization of the 120 assembled viral genomes (Fig. 2a and Fig. S1). Most viruses encoded a single protein. For example, all viruses in the families *Botourmiaviridae*, *Narnaviridae*, *Partitiviridae*, *Bromoviridae*, *Circoviridae*, *Phasmaviridae*, and *Rhabdoviridae* encoded an RdRp. Out of nine viruses in the family *Gammaflexiviridae* encoding an RdRp, three viruses encoded RdRp and movement proteins (MPs). In the family *Totiviridae*, many identified viruses had both coat proteins (CPs) and RdRp proteins or a single CP or RdRp, except for the genome of Uromyces totivirus D, which encoded two hypothetical proteins. Some viruses in the families *Potyviridae*, *Fusariviridae*, *Secoviridae*, *Hypoviridae*, and *Tymoviridae* encoded a polyprotein. The genome organization for viruses in the family *Virgaviridae* and unclassified viruses was very diverse. For instance, Entomophthora virgavirus A encoded a polyprotein and three hypothetical proteins, whereas Leucocoprinus tobamovirus A encoded an RdRp and MP.

**Figure 2.**
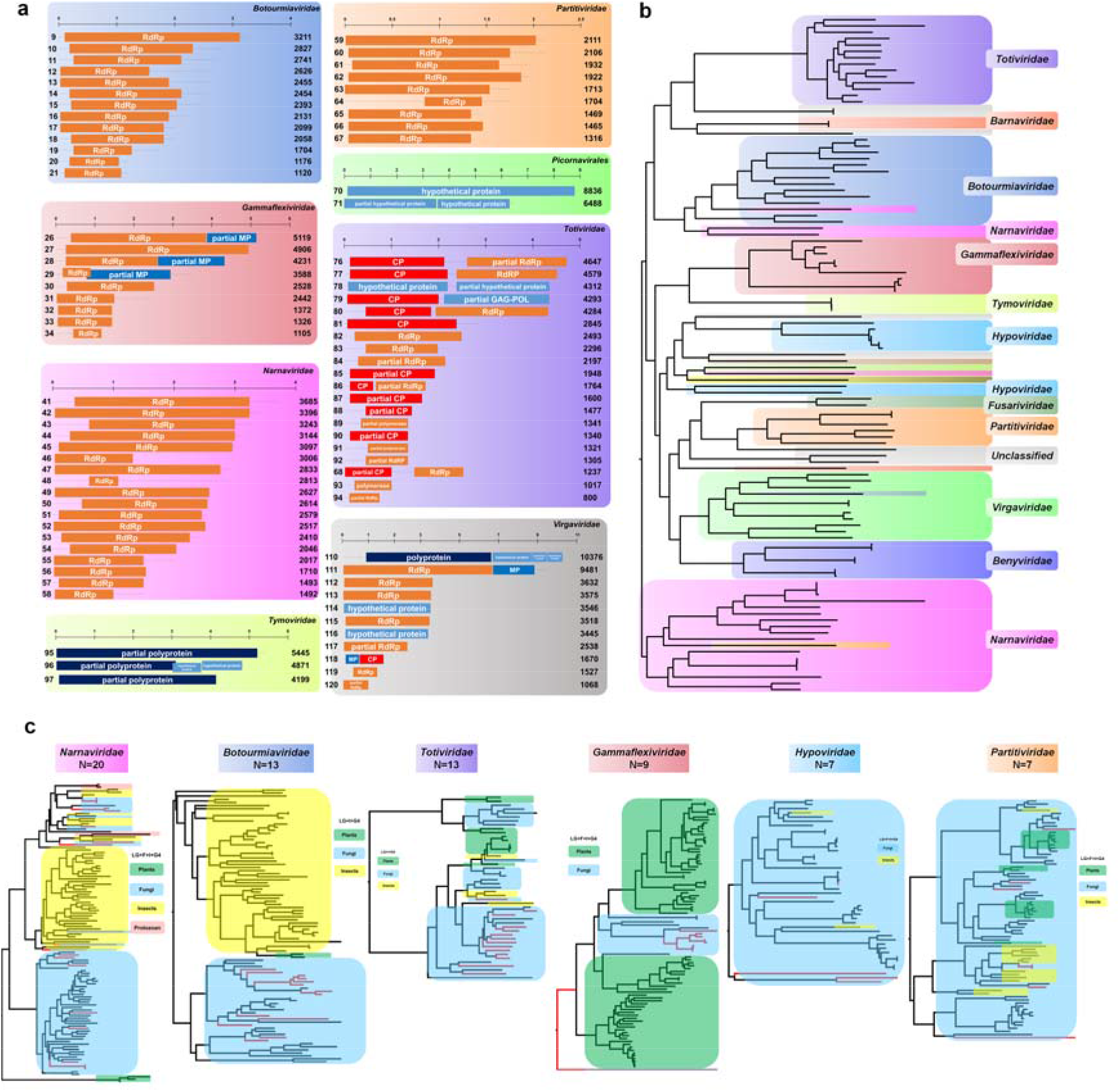
Phylogenetic trees and genome organization for assembled viruses. (a) Genome organization for identified viruses in seven viral families and one viral order. Each virus protein was visualized by different colored boxes. The number on the left in each virus corresponds to the number in Table S2. The number on the right indicates virus genome size. Genome organization for viruses in other families can be found in Fig. S2. (b) Phylogenetic tree using RdRp sequences of 107 identified viruses. Amino acid sequences were aligned with MAFFT and used for phylogenetic tree construction using maximum likelihood method and the LG+I+G4 protein substitution model implemented in IQ-TREE. The phylogenetic tree is midpoint rooted using FigTree. Viruses were indicated by different colors according to corresponding virus family. The detailed phylogenetic tree can be found in Fig. S2. (c) Phylogenetic trees of identified viruses belonging to six major families and related viruses. RdRp sequences of identified viruses and related viruses were aligned by MAFFT followed by sequence trimming using TrimAL. Maximum likelihood phylogenetic trees for individual virus families were constructed using IQ-TREE. The protein substitution model was indicated. Red lines indicate viruses identified from our study. Viruses from plants, fungi, and insects were indicated by different colored boxes. The detailed phylogenetic trees can be found in Figs. S3–S8.

Based on 107 RdRp amino acid sequences, we generated a phylogenetic tree revealing 16 viral families, one viral order, and one group with unclassified viruses (Fig. 2b and Fig. S2). To reveal the phylogenetic relationships of the identified viruses in detail, we generated 18 phylogenetic trees according to virus families (Figs. S3–S20). The six major virus families identified were *Narnaviridae* (20 viruses), *Botourmiaviridae* (13 viruses), *Totiviridae* (13 viruses), *Gammaflexiviridae* (nine viruses), *Hypoviridae* (seven viruses), and *Partitiviridae* (seven viruses) (Fig. 2c). The phylogenetic trees revealed the genetic relationships and host interactions of the identified viruses (Fig. 3c). For instance, all viruses in the families *Botourmiaviridae*, *Gammaflexiviridae*, and *Hypoviridae* as well as nine mitoviruses in the family *Narnaviridae* derived from this study were closely related to those from fungal hosts. However, 11 narnaviruses, a totivirus, and two partitiviruses were grouped with those from insects.

**Figure 3.**
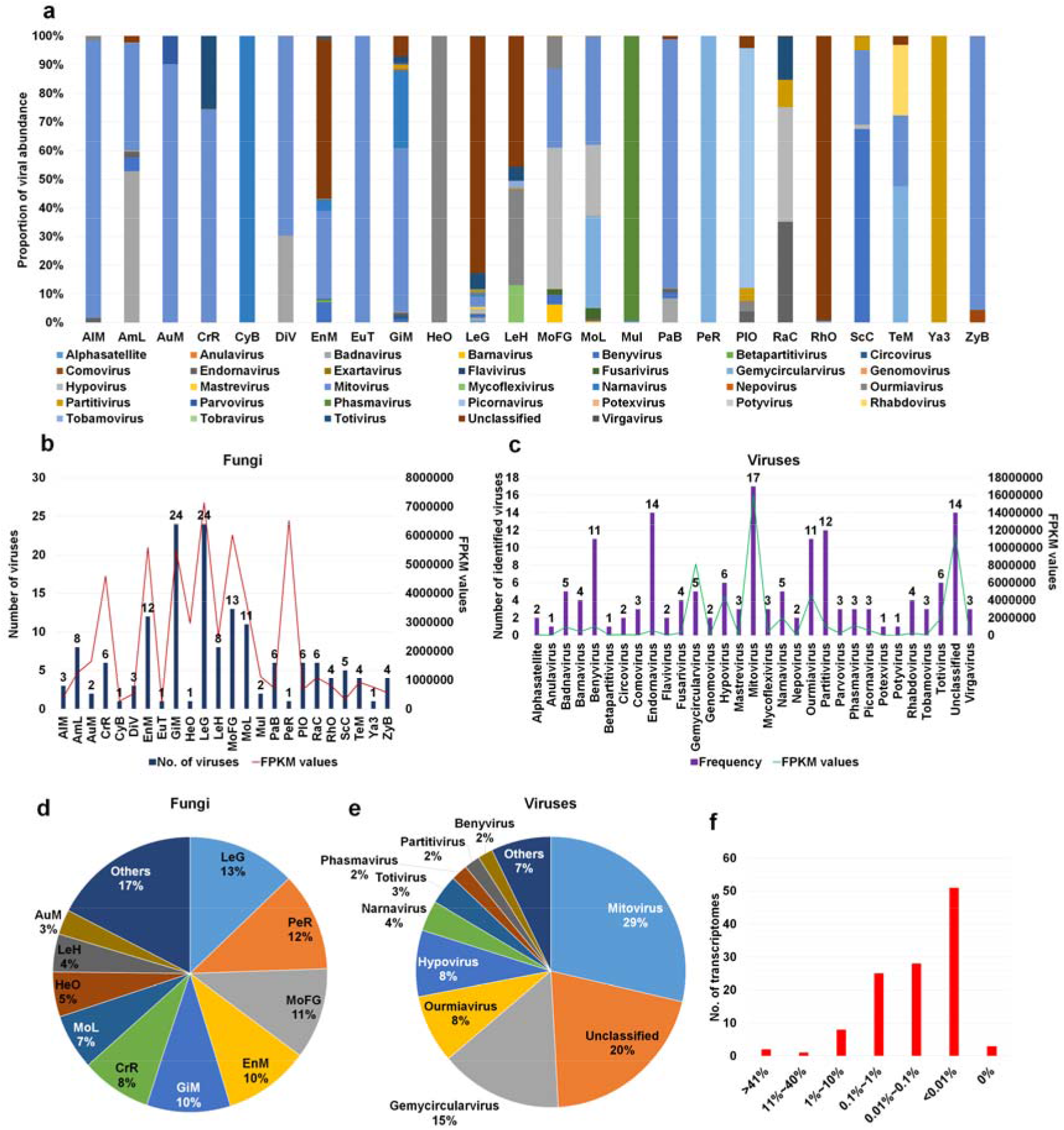
Number of identified viruses and viral abundance in individual fungal species. (a) Proportion of identified viruses based on viral abundance. Viral abundance of each virus in identified fungal transcriptomes was calculated by FPKM values. FPKM values for identified viruses in same fungal species were combined for simplicity. (b) Number of identified viruses and total viral abundance in each fungal species. (c) Number of viruses infecting fungal transcriptomes for individual virus species and total viral abundance of individual virus species. (d) Proportion of fungal species according to viral abundance. (e) Proportion of identified viruses according to viral abundance. (f) Number of fungal transcriptomes according to viral abundance. Viral abundance in each fungal transcriptome was measured by number of virus-associated reads divided by total reads.

The 20 viruses identified in the family *Narnaviridae* were further divided into narnaviruses (11 viruses) and mitoviruses (nine viruses) (Fig. S3). Most narnaviruses were closely related to known narnaviruses from insects except for two Puccinia narnaviruses, which were grouped with the Fusarium poae narnavirus 1. Interestingly, four narnaviruses derived from trypanosomatids, which are parasites of insects, were clustered together with those from insects and fungi. We identified 13 ourmiaviruses in the family *Botourmiaviridae* from six fungal species in the same clade, which were distantly grouped with two ourmiaviruses from plants (Fig. S4). Interestingly, four different *Monilinia* ourmiaviruses were phylogenetically different from each other. Except for Uromyces totivirus D in the same clade with two Wuhan insect viruses, 13 totiviruses from *Uromyces*, *Podosphaera*, and *Phakopsora* genera were together with other totiviruses from fungi (Fig. S5). Of nine gammaflexiviruses from *Leucocoprinus* species, Leucocoprinus gammaflexivirus E was distantly related with other gammaflexiviruses (Fig. S6). Except for Mycosphaerella hypovirus A, six hypoviruses were identified from *Monilinia* species (Fig. S7). The phylogenetic tree showed that all known hypoviruses were identified from fungi except for two hypoviruses from insects. Among the seven identified partitiviruses, Entomophthora partitivirus D was closely related to other partitiviruses from insects (Fig. S8).

Two identified barnaviruses in this study were very distantly related with known Rhizoctonia solani barnavirus 1 (Fig. S9). Out of six benyviruses, we identified two Hubei Beny-like viruses from *Entomophthora muscae* causing a fungal disease in flies, while the other four viruses were grouped with those from fungi (Fig. S10). We identified an anulavirus in the family *Bromoviridae*, a circovirus in the family *Circoviridae*, and two fusariviruses from fungi (Figs. S11–S13). The Mucor phasmavirus A in this study was grouped with Mucorales RNA virus 1 and two other insect viruses (Fig. S14). Of two picornaviruses, the novel Pleurotus picornavirus A from mushroom showed strong similarity to known Dicistroviridae TZ-1 in the family *Dicistroviridae* from human blood (Fig. S15). Uromyces potyvirus A and Zymoseptoria comovirus A were grouped with other viruses from plants (Fig. S16 and Fig. S17). Entomophthora rhabdovirus A was closely related with other insect rhabdoviruses (Fig. S18). Three tymoviruses from *Leucocoprinus* species and all six unclassified viruses were grouped with those from fungi (Fig. S19 and Fig. S20).

### Viral abundance in different fungal transcriptomes

We examined viral abundance in each fungal transcriptome based on FPKM (fragments per kilobase of exon model per million reads mapped) values by mapping raw sequence reads from SRA data on the 468 virus-associated contigs. Unfortunately, few SRA datasets were available (Table S4). After calculating FPKM values for each SRA dataset, we found that the SRA data from the FLX454 system were highly mapped on the virus-associated contigs due to long read sizes (Table S5). Therefore, we excluded seven SRA datasets from further analysis. Finally, we obtained FPKM values from 111 SRA datasets (Table S5). To simplify the process, we combined the viral FPKM values for each identified virus species in the same fungal species (Table S6). The 24 fungal species showed unique viral populations (Fig. 3a). Mitoviruses were dominant viruses in transcriptomes of *Alternaria* species (AlM), *Aureobasidium melanogenum* (AuM), *Cronartium ribicola* (CrR), *Eutiarosporella tritici-australis* (EuT), and *Paraglomus brasilianum* (PaB). In some fungal species, a single virus was identified, such as narnavirus in *Cyathus bulleri* (CyB), phasmavirus in *Mucor irregularis* (Mul), gemycircularvirus in *Pecoramyces ruminatium* (PeR), unclassified virus in *Rhizopus oryzae* (RhO), and partitivirus in *Yarrowia* species (Ya3). *Leucoagaricus gongylophorus* (LeG) contained several viruses; however, an unclassified virus was dominantly present. *Racocetra castanea* (RaC) contained four different viruses, and of them, endornavirus and hypovirus were dominantly present.

We compared the number of infecting viruses and the amount of viral abundance (Fig. 3b) and found that the number of infecting viruses was correlated with viral abundance. For example, both GiM and LeG contained 24 different viruses with a high amount of viral abundance. However, PeR was infected by a single virus, but its viral abundance was very high. Next, we examined the frequency of identified viruses in different fungal species (Fig. 3c). Anulavirus, betapartitivirus, potexvirus, and potyvirus were identified in a single fungal species, whereas mitovirus was identified in at least 17 fungal species. Moreover, endornavirus (14 species), the unclassified virus (14 species), partitivirus (12 species), barnavirus (11 species), and ourmiavirus (11 species) were frequently identified in different fungal species. Based on FPKM values, mitovirus showed the highest viral abundance followed by the unclassified virus and gemycircularvirus.

After combining virus-associated FPKM values, we examined the proportion of identified viruses within individual fungal species (Fig. 3d). The proportions of virus-associated FPKM values were very high in LeG (13%), PeR (12%), and MoFG (11%). Based on FPKM values, mitovirus (29%) was the most abundantly present virus followed by the unclassified virus (20%), gemycircularvirus (15%), ourmiavirus (8%), and hypovirus (8%) (Fig. 3e). Finally, we examined the proportion of mycoviruses in each fungal transcriptome based on sequence reads (Fig. 3f and Table S7). The proportion of virus-associated reads in most transcriptomes ranged from 0.01% to 0.1%; however, the proportion of virus-associated reads in two transcriptomes of *Leucoagaricus gongylophorus* was higher than 41% (Table S7).

### Analyses of diverse *Monilinia* and *Gigaspora* viromes

We examined changes of viromes for two selected fungal transcriptome projects for *Monilinia* and *Gigaspora* species, respectively. *Monilinia* transcriptomes comprised 18 different samples established from three different hosts, four different regions, and three different *Monilinia* species: *M. fructicola*, *M. laxa*, and *M. fructigena* (36). The number of identified viruses ranged from one to 10 (Table S8). In particular, three samples from *M. fructigena* collected from pear in Emilia-Romagna were severely infected by many viruses (nine to 10 viruses) (Fig. 4a). In addition, the proportion of virus-associated reads in three *M. fructigena* transcriptomes was relatively high (2.3–6.1%) as compared to those of the other two *Monilinia* species (less than 1%).

**Figure 4.**
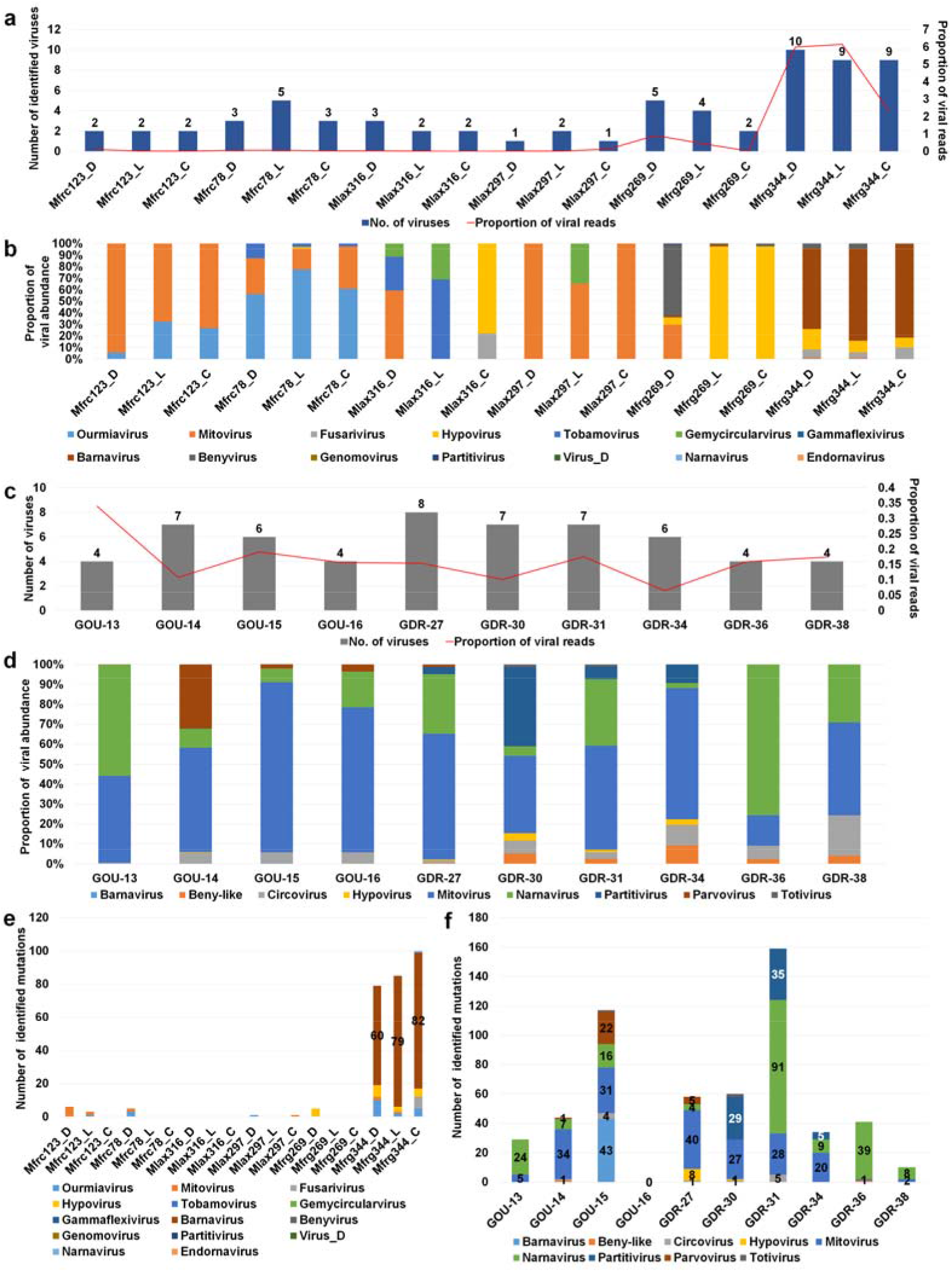
Analyses of viromes for three different *Monilinia* species and *Gigaspora margarita*. (a) Number of identified viruses (left) and proportion of viral abundance (%) (right) in 18 different *Monilinia* transcriptomes. (b) Proportion of identified viruses in each *Monilinia* transcriptome. (c) Number of identified viruses (left) and proportion of viral abundance (%) (right) in 10 different *G. margarita* transcriptomes. (d) Proportion of identified viruses in each *G. margarita* transcriptome. Number of identified virus mutations for individual virus species in each *Monilinia* (e) and *G. margarita* (f) transcriptome.

We examined the proportion of infecting viruses in each *Monilinia* transcriptome (Fig. 4b). In general, the samples from the same host and region showed similar viral populations except for three *M. laxa* samples derived from cherry in Puglia showing very different viral populations. *M. fructicola* collected from cherry in two different regions showed region-specific viral populations. For example, mitovirus was dominant in *M. fructicola* from the Puglia region, while ourmiavirus was dominant in *M. fructicola* from the Campania region. Although different growth conditions and tissues were used, we did not find any possible effects on viral populations. Hypovirus was dominant in two samples from plum in the Basilicata region, whereas barnavirus was dominant in three samples from pear in the Emilia-Romagna region.

Next, we analyzed viromes of *Gigaspora margarita*, which comprised 21 transcriptomes derived from 10 different conditions (Table S9) (37). *G. margarita* was infected by at least nine different viruses, and the proportion of virus-associated reads ranged from 0.06–0.33% (Fig. 4c). The GOU-13 sample was infected by only four viruses; however, the proportion of virus-associated reads was the highest (0.33%). Two frequently identified viruses in *G. margarita* were mitovirus and narnavirus (Fig. 4d). In particular, mitovirus was the most dominant virus in six out of 10 conditions. Although *G. margarita* was derived from a single strain, the viral populations in each condition were varied. Strigolactone is known to stimulate arbuscular mycorrhizal fungi by activating mitochondria (37). GOU-15, GDR-31, and GDR-34 samples were treated with strigolactone; however, we did not observe any significant changes in the viromes of *G. margarita* caused by strigolactone. This study was initially conducted to observe the symbiotic effects of an endobacterium to increase the bioenergetic potential of *G. margarita* (37). The number of infecting viruses and viral abundance were slightly reduced in three symbiotic conditions—GDR-16, GDR-36, and GDR-38—as compared to the other seven pre-symbiotic conditions.

Viruses have high mutation rates as compared to their hosts. We examined virus mutation frequency in two different viromes. Although *Monilinia* was infected by many viruses, only three samples from *M. fructigena* collected from pear in Emilia-Romagna showed many mutations (Table S10). In particular, barnavirus showed many mutations as compared to other viruses (Fig. 4e). By contrast, several mutations were observed in most *G. margarita* samples (Fig. 4f and Table S11). Interestingly, GOU15 and GDR-31 samples treated with strigolactone had many mutations in several identified virus genomes. In GOU-15 containing six viruses, barnavirus (43 mutations) had the highest number of mutations followed by mitovirus (31 mutations) and parvovirus (22 mutations). In GDR-31 containing four viruses having mutations, narnavirus (91 mutations) had the highest number of mutations followed by partitivirus (35 mutations) and narnavirus (28 mutations). Interestingly, the numbers of virus mutations for two symbiotic conditions, GOU-16 and GOU-38, were zero and 10, respectively.

## DISCUSSION

Here, we conducted a large-scale *in silico* mycovirus identification and mycovirome studies using public fungal transcriptome datasets. Our study focused on mycoviruses, which have not been well explored as compared to viruses infecting other eukaryotic kingdoms. Similarly, a recent study also carried out *in silico* bioinformatics analyses using available fungal transcriptomes identifying 59 mycoviral genomes (21). There were two main differences in materials and methods between two studies. The first was the diversity of fungal transcriptomes. In our study, we analyzed 126 fungal transcriptomes from at least 11 fungal phyla whereas the recent study focused on Pezizomycotina fungi of the phylum *Ascomycota*. The second was reference viral proteins for virus identification. To increase the number of virus-associated contigs, we used all available viral proteins for the BLASTX search instead of using viral RdRp sequences, revealing a wide range of mycoviruses encoding diverse viral proteins. By contrast, the recent study only used six different viral RdRp families. Our approach enabled us assemble 120 viral genomes and reveal mycoviruses classified into five orders, 21 families, 26 genera, and 88 species, representing the widest range of mycoviral taxonomy in a single study. Although both studies analyzed assembled mycoviral genomes with phylogenetic trees and domain study, our study additionally showed viral abundance, viral diversity, and mutation frequency using fungal transcriptomes. Our results might provide a better picture of how much of the true overall mycovirome is visible within these results.

To increase specificity for virus identification, BLASTX against the NR database eliminated possible endogenous virus-like sequences, resulting in 468 virus-associated contigs. Due to the low abundance of mycoviruses in fungal transcriptomes, the number of mycovirus-associated reads was very low resulting that about half of the assembled virus-associated contigs were less than 1 kb. Moreover, it is also possible that 83 of 126 fungal transcriptomes that did not contain any virus-associated contigs might be infected by unknown mycoviruses. The low abundance of mycoviruses in the fungal hosts could not be identified by metatranscriptomics; however, they could be identified PCR based approaches with virus-specific primers. Identification of virus by a single approach does not guarantee virus infection in a living organism.

Most identified virus-associated contigs were derived from ssRNA and dsRNA genomes; however, we identified viruses with negative ssRNA, ssDNA, and dsDNA genomes, suggesting transcribed viral RNAs from DNA genomes by metatranscriptomics. For example, the infection of gemycircularviruses and genomoviruses with a single circular DNA genome in fungi has been reported previously (18); however, this is the first report of identification of a mastrevirus and alphasatellite from fungal transcriptomes.

The phylogenetic tree revealed that many of the identified mycoviruses were originated from fungi; however, some mycoviruses had strong phylogenetic relationships with those from insects and plants. In particular, four known narnaviruses derived from trypanosomatids were closely related with those from insects and fungi, suggesting that the previously identified narnaviruses from insects might be derived from insect parasites (38). It is likely that the insect parasites were also included during RNA preparation from insect tissues.

Several virus families, including *Bromoviridae*, *Circoviridae*, and *Dicistroviridae*, have been identified from fungi for the first time. Interestingly, Gigaspora circovirus A with ssDNA genomes showed sequence similarity to only beak and feather disease virus, which causes a viral disease in birds (39). Circoviruses are circular ssDNAs that encode for two proteins bidirectional replication initiator protein (Rep) and CP. However, Gigaspora circovirus A has only a Rep. Furthermore, Gigaspora circovirus A was identified in all 10 *Gigaspora* transcriptomes. This result confirmed the possible infection of a circovirus in the fungus. The strong similarity between the novel Pleurotus picornavirus A and known Dicistroviridae TZ-1 from the blood of febrile Tanzanian children was somewhat interesting (40). However, other dicistroviruses from insects were grouped together with picornaviruses from insects, suggesting the presence of dicistrovirus in the mushroom. In addition, parvoviruses and flaviviruses, which infect both vertebrate and invertebrate hosts, were detected in fungi for the first time.

Uromyces potyvirus A and Zymoseptoria comovirus A showed strong similarity to other viruses from plants, indicating that both fungi transmit plant viruses that infect both fungi and plants. Fungal transmission of a plant virus has been reported (41). Moreover, plants are known hosts for viruses in the family *Virgaviridae*, which were also identified in fungi, suggesting that fungi could be alternative hosts. A previous study reporting the replication of endophytic mycovirus in plant cells supports our finding (42).

The genome organization of the 120 mycoviromes indicates that most mycoviruses had simple genome structures. For instance, many of the identified mycoviruses had an RdRp required for viral replication. In addition, we found that the genome structures of some viruses in the families *Barnaviridae*, *Benyviridae*, *Gammaflexiviridae*, *Virgaviridae*, and *Tymoviridae* were not identical, indicating changes of viral genome structures during the mycovirus’s evolution and host adaptation.

A few recent studies revealed the complete genome of a large number of viruses from vertebrates and invertebrates using transcriptomic and complementary approaches (3,4). By contrast, in our *in silico* mycovirome study, we faced several obstacles to obtaining complete mycoviral genomes from only transcriptome data because of the low titer of mycoviruses in the fungi and unavailability of original samples. However, 468 mycovirus-associated sequences, in which 120 viral genomes were assembled, provided a comprehensive overview of the distribution of mycovirus taxonomy and their hosts. It was also surprising that the fungi could be the hosts of numerous viruses, as shown in GiM and LeG, which were infected by at least 24 different viruses. In addition, individual mycoviromes showed at least 10 and eight different viruses infecting *Monilinia* and *Gigaspora* species, respectively, revealing the coinfection of mycoviruses in many fungal hosts.

The virome study based on the mapping of raw sequence reads revealed a possible correlation between the number of infecting viruses and viral abundance. This result suggests that the competition of diverse coinfected mycoviruses promoted the viral replication. Although the coinfection of diverse mycoviruses in the fungi was common in our study, most viromes had a dominant virus species. Our study identified the most common mycoviruses, such as mitovirus, endornavirus, benyvirus, ourmiavirus, and partitivirus, with a wide range of fungal hosts, as described previously (20,21). However, several viruses, including anulavirus, betapartitvirus, potexvirus, and potyvirus, had a limited range of hosts.

The single nucleotide polymorphism (SNP) analysis showed that there were differences in virus mutation frequency among coinfecting mycoviruses, and most mycoviruses did not show strong mutation except for a few mycoviruses, such as barnavirus, mitovirus, and narnavirus.

The study of *Monilinia* viromes revealed that there were strong deviations in the composition and viral abundance of viruses that originated from identical samples. This might have been caused by virus abundance, RNA extraction, library preparation, or RNA-seq. In addition, hosts and sample regions played an important role in viral populations in different *Monilinia* species. Although different dark and light conditions and tissues were used for sample preparation, it was not sufficient to determine their effects on the virome without biological replicates.

The study of *Gigaspora* viromes revealed that the chemical strigolactone might promote virus replication and mutations, while symbiosis with endobacteria might suppress virus replication and mutations. Strigolactones, plant hormones extracted from plant roots, play an important role in the symbiotic interaction between plants and arbuscular mycorrhizal fungi. However, our results demonstrated negative effects of strigolactones on fungi and mycoviruses. The strong symbiotic interaction between endobacteria and *Gigaspora* species inhibited mycovirus replication. Thus, the effects of strigolactones and the symbiosis of endobacteria on mycovirus lifecycles should be further elucidated.

As shown in our study, the compositions and populations of mycoviruses were more complex than we expected. This study is the largest comprehensive mycovirome study shedding light on an array of mycovirome-associated topics, including virus identification, fungal host ranges, taxonomy of mycoviruses, viral genome assembly, phylogenetic analyses, viral abundance, mutations, and changes of mycoviral populations in different conditions. Moreover, this study provides many clues and information to reveal the diversity and host distribution of mycoviruses.

## DATA AVAILABILITY

Viral genome sequences for 120 mycoviruses were deposited in GenBank with the following accession numbers: MK231015–MK231133 and MK940812.

## SUPPLEMENTARY DATA

### Supplementary Figures

Figure S1. Genome organization for identified viruses in families *Barnaviridae*, *Benyviridae*, *Bromoviridae*, *Circoviridae*, *Fusariviridae*, *Hypoviridae*, *Phasmaviridae*, *Potyviridae*, *Rhabdoviridae*, and *Secoviridae* and unclassified viruses.

Figure S2. Phylogenetic tree of 107 identified viruses using RdRp sequences.

Figure S3. Phylogenetic tree of 20 viruses in family *Narnaviridae* and related viruses.

Figure S4. Phylogenetic tree of 13 viruses in family *Botourmiaviridae* and related viruses.

Figure S5. Phylogenetic tree of 14 viruses in family *Totiviridae* and related viruses.

Figure S6. Phylogenetic tree of nine viruses in family *Gammaflexiviridae* and related viruses.

Figure S7. Phylogenetic tree of seven viruses in family *Hypoviridae* and related viruses.

Figure S8. Phylogenetic tree of seven viruses in family *Partitiviridae* and related viruses.

Figure S9. Phylogenetic tree of two viruses in family *Barnaviridae* and related viruses.

Figure S10. Phylogenetic tree of six viruses in family *Benyviridae* and related viruses.

Figure S11. Phylogenetic tree of a virus in family *Bromoviridae* and related viruses.

Figure S12. Phylogenetic tree of a virus in family *Circoviridae* and related viruses.

Figure S13. Phylogenetic tree of two viruses in family *Fusariviridae* and related viruses.

Figure S14. Phylogenetic tree of a virus in the family *Phasmaviridae* and related viruses.

Figure S15. Phylogenetic tree of two viruses in the order *Picornavirales* and related viruses.

Figure S16. Phylogenetic tree of a virus in the family *Potyviridae* and related viruses.

Figure S17. Phylogenetic tree of a virus in the family *Secoviridae* and related viruses.

Figure S18. Phylogenetic tree of a virus in the family *Rhabdoviridae* and related viruses.

Figure S19. Phylogenetic tree of three viruses in the family *Tymoviridae* and related viruses.

Figure S20. Phylogenetic tree of six unclassified viruses and related viruses.

### Supplementary Tables

Table S1. Summary of fungal transcriptomes used for virus identification.

Table S2. Summary of BLASTX results for 468 virus-associated contigs against viral proteins.

Table S3. Detailed information of 120 viral genomes assembled from fungal transcriptomes in this study.

Table S4. Information of SRA data used for virome study.

Table S5. Mapping results of raw sequence reads for each SRA dataset on 468 virus-associated contigs.

Table S6. FPKM values for identified viruses in each fungal species.

Table S7. Mapping results of raw sequence reads for each SRA dataset on 468 virus-associated contigs.

Table S8. Detailed information for *Monilinia* viromes.

Table S9. Detailed information for *Gigaspora margarita* viromes.

Table S10. Number of identified mutations for viruses identified from *Monilinia* transcriptomes.

Table S11. Number of identified mutations for viruses identified from *Monilinia* transcriptomes.

## ACKNOWLEDGEMENTS

I (Won Kyong Cho) would like to express our deepest gratitude to Hyang Sook Kim, Mi Kyong Kim, and Jin Kyong Cho for all their support with the current project. This work is dedicated to the memory of my father, Tae Jin Cho (1946–2015).

## REFERENCES

1. Lefkowitz, E.J., Dempsey, D.M., Hendrickson, R.C., Orton, R.J., Siddell, S.G., and Smith, D.B., (2017) Virus taxonomy: the database of the International Committee on Taxonomy of Viruses (ICTV). Nucl Acid Res, 46, D708–D717.

2. Monier, A., Claverie, J.-M., and Ogata, H. (2008) Taxonomic distribution of large DNA viruses in the sea. Genome Biol, 9, R106.

3. Shi, M., Lin, X.-D., Tian, J.-H., Chen, L.-J., Chen, X., Li, C.-X., Qin, X.-C., Li, J., Cao, J.-P., and Eden, J.-S., (2016) Redefining the invertebrate RNA virosphere. Nature, 540, 539.

4. Shi, M., Lin, X.-D., Chen, X., Tian, J.-H., Chen, L.-J., Li, K., Wang, W., Eden, J.-S., Shen, J.-J., and Liu, L. (2018) The evolutionary history of vertebrate RNA viruses. Nature, 556, 197.

5. Mushegian, A., Shipunov, A. and Elena, S.F., (2016) Changes in the composition of the RNA virome mark evolutionary transitions in green plants. BMC Biol, 14, 68.

6. Shkoporov, A.N., and Hill, C. (2019) Bacteriophages of the human gut: the “known unknown” of the microbiome. Cell Host Microbe, 25, 195–209.

7. Prussin, A.J., Marr, L.C., and Bibby, K.J., (2014) Challenges of studying viral aerosol metagenomics and communities in comparison with bacterial and fungal aerosols. FEMS Microbiol Lett, 357, 1–9.

8. Kumar, N., Barua, S., Riyesh, T., Chaubey, K.K., Rawat, K.D., Khandelwal, N., Mishra, A.K., Sharma, N., Chandel, S.S., and Sharma, S. (2016) Complexities in isolation and purification of multiple viruses from mixed viral infections: viral interference, persistence and exclusion. PLoS One, 11, e0156110.

9. Nowrousian, M. (2010) Next-generation sequencing techniques for eukaryotic microorganisms: sequencing-based solutions to biological problems. Eukaryotic Cell, 9, 1300–1310.

10. Zhang, Y.-Z., Shi, M. and Holmes, E.C., (2018) Using metagenomics to characterize an expanding virosphere. Cell, 172, 1168–1172.

11. Ho, T. and Tzanetakis, I.E., (2014) Development of a virus detection and discovery pipeline using next generation sequencing. Virology, 471, 54–60.

12. Yamashita, A., Sekizuka, T. and Kuroda, M. (2016) VirusTAP: viral genome-targeted assembly pipeline. Front Microbiol, 7, 32.

13. Prosperi, M.C., Prosperi, L., Bruselles, A., Abbate, I., Rozera, G., Vincenti, D., Solmone, M.C., Capobianchi, M.R., and Ulivi, G. (2011) Combinatorial analysis and algorithms for quasispecies reconstruction using next-generation sequencing. BMC Bioinform, 12, 5.

14. Watson, S.J., Welkers, M.R., Depledge, D.P., Coulter, E., Breuer, J.M., de Jong, M.D., and Kellam, P. (2013) Viral population analysis and minority-variant detection using short read next-generation sequencing. Philos Trans Royal Soc B, 368, 20120205.

15. Paez-Espino, D., Eloe-Fadrosh, E.A., Pavlopoulos, G.A., Thomas, A.D., Huntemann, M., Mikhailova, N., Rubin, E., Ivanova, N.N., and Kyrpides, N.C., (2016) Uncovering Earth’s virome. Nature, 536, 425.

16. Nuss, D.L., (2005) Hypovirulence: mycoviruses at the fungal–plant interface. Nat Rev Microbiol, 3, 632.

17. Liu, L., Xie, J., Cheng, J., Fu, Y., Li, G., Yi, X. and Jiang, D. (2014) Fungal negative-stranded RNA virus that is related to bornaviruses and nyaviruses. Proc Nat Acad Sci, 111, 12205–12210.

18. Yu, X., Li, B., Fu, Y., Jiang, D., Ghabrial, S.A., Li, G., Peng, Y., Xie, J., Cheng, J. and Huang, J. (2010) A geminivirus-related DNA mycovirus that confers hypovirulence to a plant pathogenic fungus. Proc Nat Acad Sci, 107, 8387–8392.

19. Son, M., Yu, J. and Kim, K.-H., (2015) Five questions about mycoviruses. PLoS Pathog, 11, e1005172.

20. Marzano, S.-Y.L., Nelson, B.D., Ajayi-Oyetunde, O., Bradley, C.A., Hughes, T.J., Hartman, G.L., Eastburn, D.M., and Domier, L.L., (2016) Identification of diverse mycoviruses through metatranscriptomics characterization of the viromes of five major fungal plant pathogens. J Virol, 90, 6846–6863.

21. Gilbert, K.B., Holcomb, E.E., Allscheid, R.L., and Carrington, J.C., (2019) Hiding in plain sight: New virus genomes discovered via a systematic analysis of fungal public transcriptomes. PLoS One, 14, e0219207.

22. Thapa, V. and Roossinck, M.J., (2019) Determinants of Coinfection in the Mycoviruses. Frontiers in cellular and infection microbiology, 9, 169.

23. Jo, Y., Choi, H., Cho, J.K., Yoon, J.-Y., Choi, S.-K., and Cho, W.K., (2015) In silico approach to reveal viral populations in grapevine cultivar Tannat using transcriptome dat. Sci Rep, 5, 15841.

24. Buchfink, B., Xie, C. and Huson, D.H., (2015) Fast and sensitive protein alignment using DIAMOND. Nat Methods, 12, 59.

25. Letunic, I., Doerks, T. and Bork, P. (2011) SMART 7: recent updates to the protein domain annotation resource. Nucl Acids Res, 40, D302–D305.

26. Katoh, K. and Standley, D.M., (2013) MAFFT multiple sequence alignment software version 7: improvements in performance and usability. Mol Biol Evol, 30, 772–780.

27. Capella-Gutiérrez, S., Silla-Martínez, J.M. and Gabaldón, T. (2009) trimAl: a tool for automated alignment trimming in large-scale phylogenetic analyses. Bioinformatics, 25, 1972–1973.

28. Nguyen, L.-T., Schmidt, H.A., von Haeseler, A. and Minh, B.Q., (2014) IQ-TREE: a fast and effective stochastic algorithm for estimating maximum-likelihood phylogenies. Mol Biol Evol, 32, 268–274.

29. Hoang, D.T., Chernomor, O., Von Haeseler, A., Minh, B.Q., and Vinh, L.S., (2017) UFBoot2: improving the ultrafast bootstrap approximation. Mol Biol Evol, 35, 518–522.

30. Guindon, S., Dufayard, J.-F., Lefort, V., Anisimova, M., Hordijk, W. and Gascuel, O. (2010) New algorithms and methods to estimate maximum-likelihood phylogenies: assessing the performance of PhyML 3.0. Sys Biol, 59, 307–321.

31. Kodama, Y., Shumway, M. and Leinonen, R. (2011) The Sequence Read Archive: explosive growth of sequencing data. Nucl Acids Res, 40, D54–D56.

32. Li, H. and Durbin, R. (2010) Fast and accurate long-read alignment with Burrows– Wheeler transform. Bioinformatics, 26, 589–595.

33. Grabherr, M.G., Haas, B.J., Yassour, M., Levin, J.Z., Thompson, D.A., Amit, I., Adiconis, X., Fan, L., Raychowdhury, R. and Zeng, Q. (2011) Full-length transcriptome assembly from RNA-Seq data without a reference genome. Nat Biotech, 29, 644.

34. Li, H., Handsaker, B., Wysoker, A., Fennell, T., Ruan, J., Homer, N., Marth, G., Abecasis, G. and Durbin, R. (2009) The sequence alignment/map format and SAMtools. Bioinformatics, 25, 2078–2079.

35. Li, H. (2011) A statistical framework for SNP calling, mutation discovery, association mapping and population genetical parameter estimation from sequencing data. Bioinformatics, 27, 2987–2993.

36. Angelini, R.M.D.M., Abate, D., Rotolo, C., Gerin, D., Pollastro, S. and Faretra, F. (2018) De novo assembly and comparative transcriptome analysis of Monilinia fructicola, Monilinia laxa and Monilinia fructigena, the causal agents of brown rot on stone fruits. BMC Genomics, 19, 436.

37. Salvioli, A., Ghignone, S., Novero, M., Navazio, L., Bagnaresi, P. and Bonfante, P. (2016) Symbiosis with an endobacterium increases the fitness of a mycorrhizal fungus, raising its bioenergetic potential. ISME J, 10, 130.

38. Grybchuk, D., Kostygov, A.Y., Macedo, D.H., Votýpka, J., Lukeš, J. and Yurchenko, V. (2018) RNA Viruses in Blechomonas (Trypanosomatidae) and Evolution of Leishmaniavirus. mBio, 9, e01932–01918.

39. Niagro, F., Forsthoefel, A., Lawther, R., Kamalanathan, L., Ritchie, B., Latimer, K. and Lukert, P. (1998) Beak and feather disease virus and porcine circovirus genomes: intermediates between the geminiviruses and plant circoviruses. Arch Virol, 143, 1723–1744.

40. Cordey, S., Laubscher, F., Hartley, M.-A., Junier, T., Pérez-Rodriguez, F.J., Keitel, K., Vieille, G., Samaka, J., Mlaganile, T. and Kagoro, F. (2019) Detection of dicistroviruses RNA in blood of febrile Tanzanian children. Emerg Microbes Infect, 8, 613–623.

41. Andika, I.B., Wei, S., Cao, C., Salaipeth, L., Kondo, H. and Sun, L. (2017) Phytopathogenic fungus hosts a plant virus: A naturally occurring cross-kingdom viral infection. Proc Nat Acad Sci, 114, 12267–12272.

42. Nerva, L., Varese, G., Falk, B. and Turina, M. (2017) Mycoviruses of an endophytic fungus can replicate in plant cells: Evolutionary implications. Sci Rep, 7, 1908.

