## Supplementary Figures for "Identification of viruses from fungal transcriptomes"

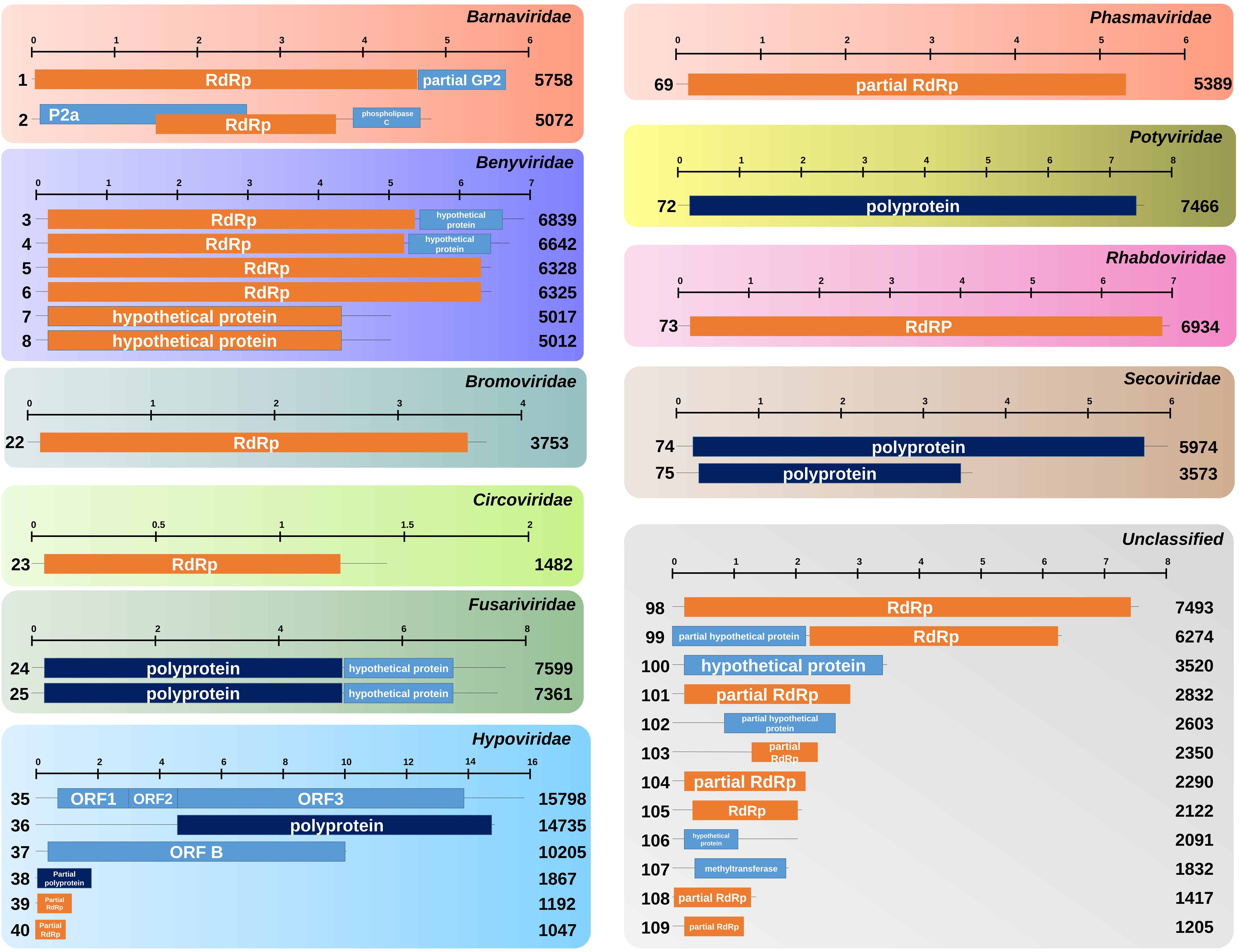


Figure S1. Genome organization for identified viruses in families *Barnaviridae*, *Benyviridae*, *Bromoviridae*, *Circoviridae*, *Fusariviridae*, *Hypoviridae*, *Phasmaviridae*, *Potyviridae*, *Rhabdoviridae*, and *Secoviridae* and unclassified viruses.

Each virus protein was visualized by different colored boxes. The number on the left in each virus corresponds to the number in Table S2. The number on the right indicates the virus genome size. Genome organization for viruses in other families can be found in Fig. 2.

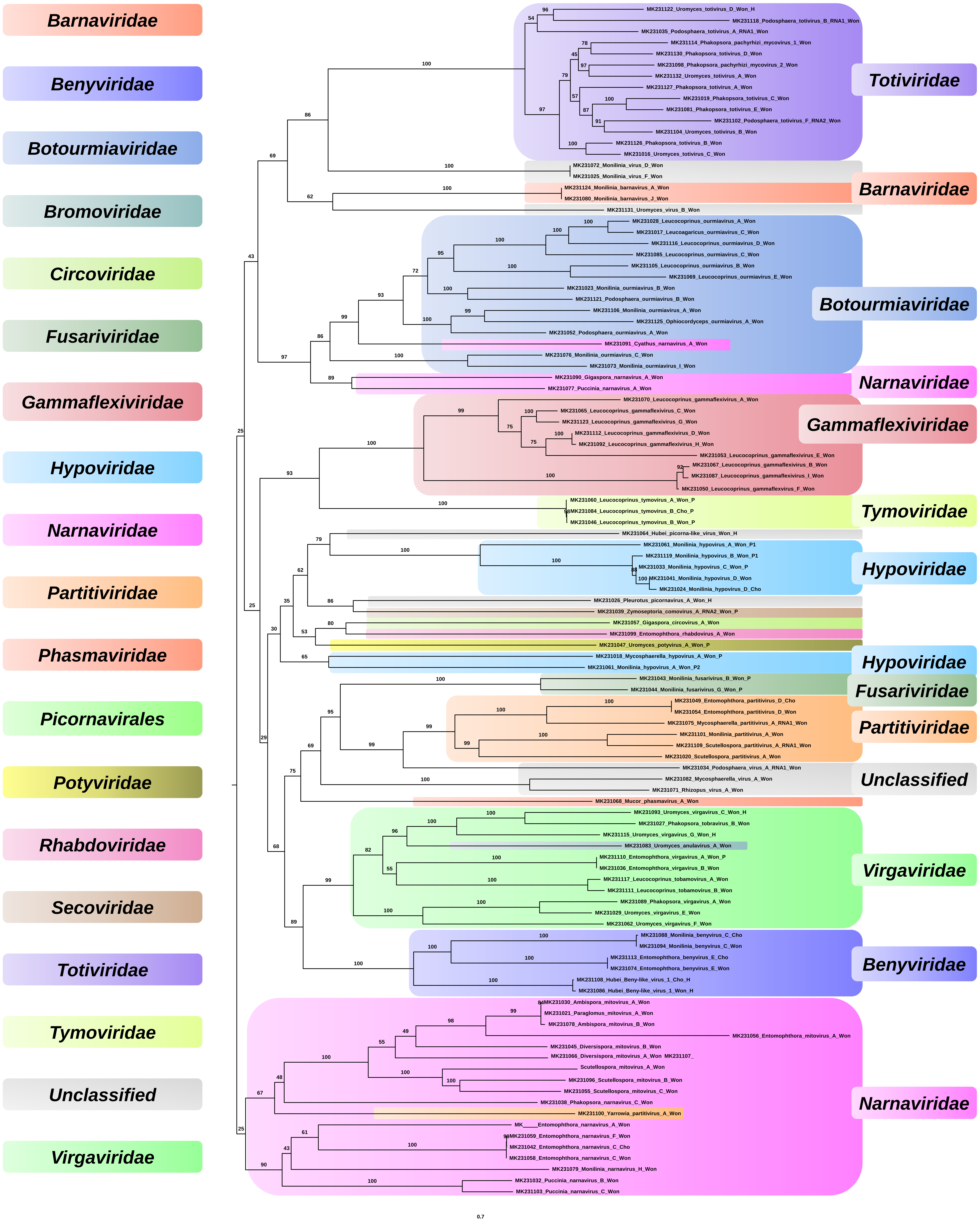


Figure S2. Phylogenetic tree of 107 identified viruses using RdRp sequences.

Amino acid sequences were aligned with MAFFT and subjected to phylogenetic tree construction using maximum likelihood method and LG+I+G4 protein substitution model implemented in IQ-TREE. The phylogenetic tree is midpoint rooted using FigTree. Viruses were indicated by different colors according to corresponding virus family. Bootstrap values are indicated on branches with 1,000 replicates.

### 20 viruses in the family *Narnaviridae*

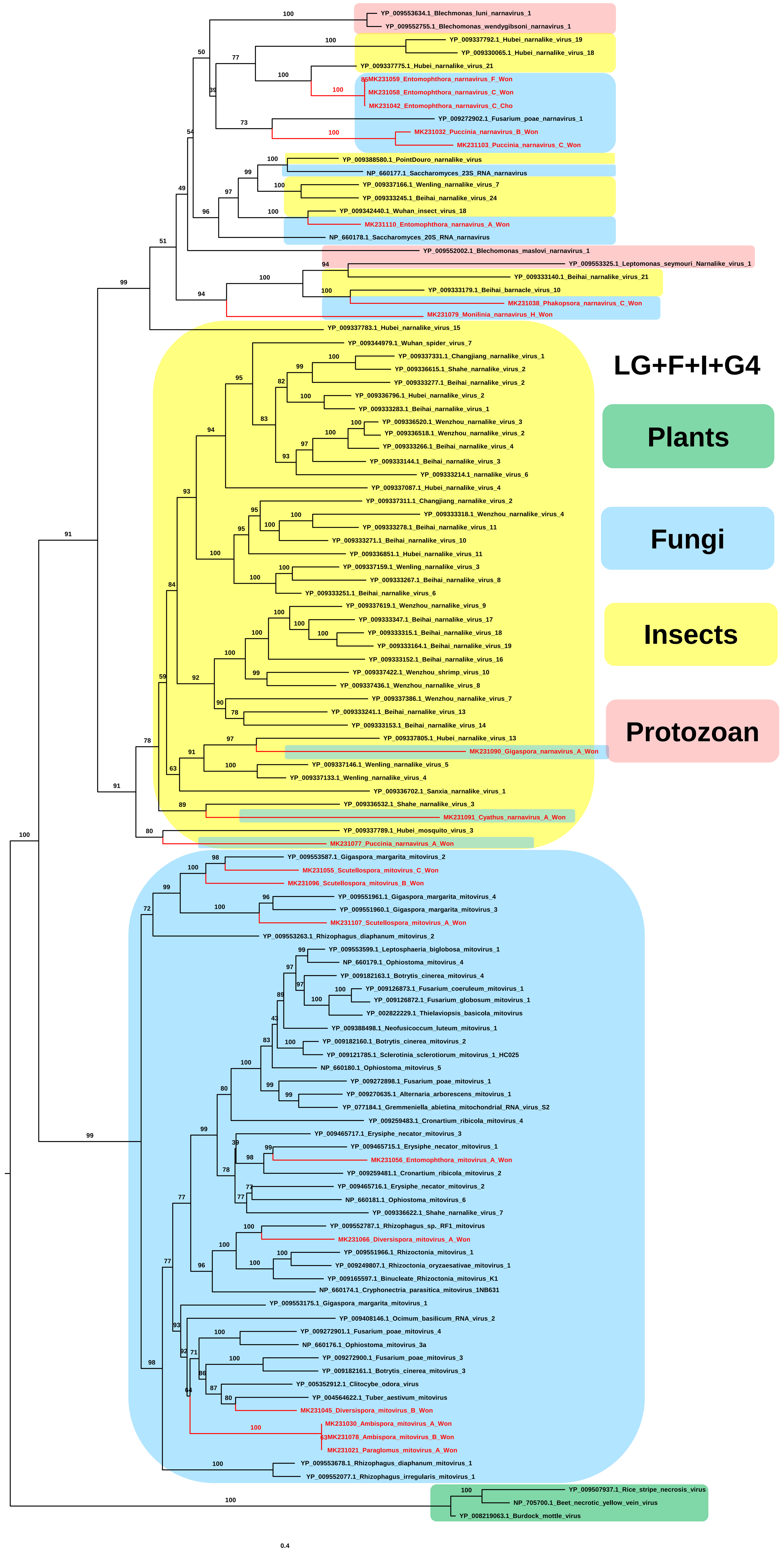


Figure S3. Phylogenetic tree of 20 viruses in family *Narnaviridae* and related viruses.
RdRp sequences for 20 viruses in family *Narnaviridae* and related viruses were aligned with MAFFT and subjected to phylogenetic tree construction using maximum likelihood method and LG+F+I+G4 protein substitution model implemented in IQ-TREE. The phylogenetic tree is midpoint rooted using FigTree. Viruses in this study were indicated by red colors. Bootstrap values are indicated on branches with 1,000 replicates. Viruses derived from different hosts were indicated by different colored boxes.

#### 13 viruses in the family *Botourmiaviridae*

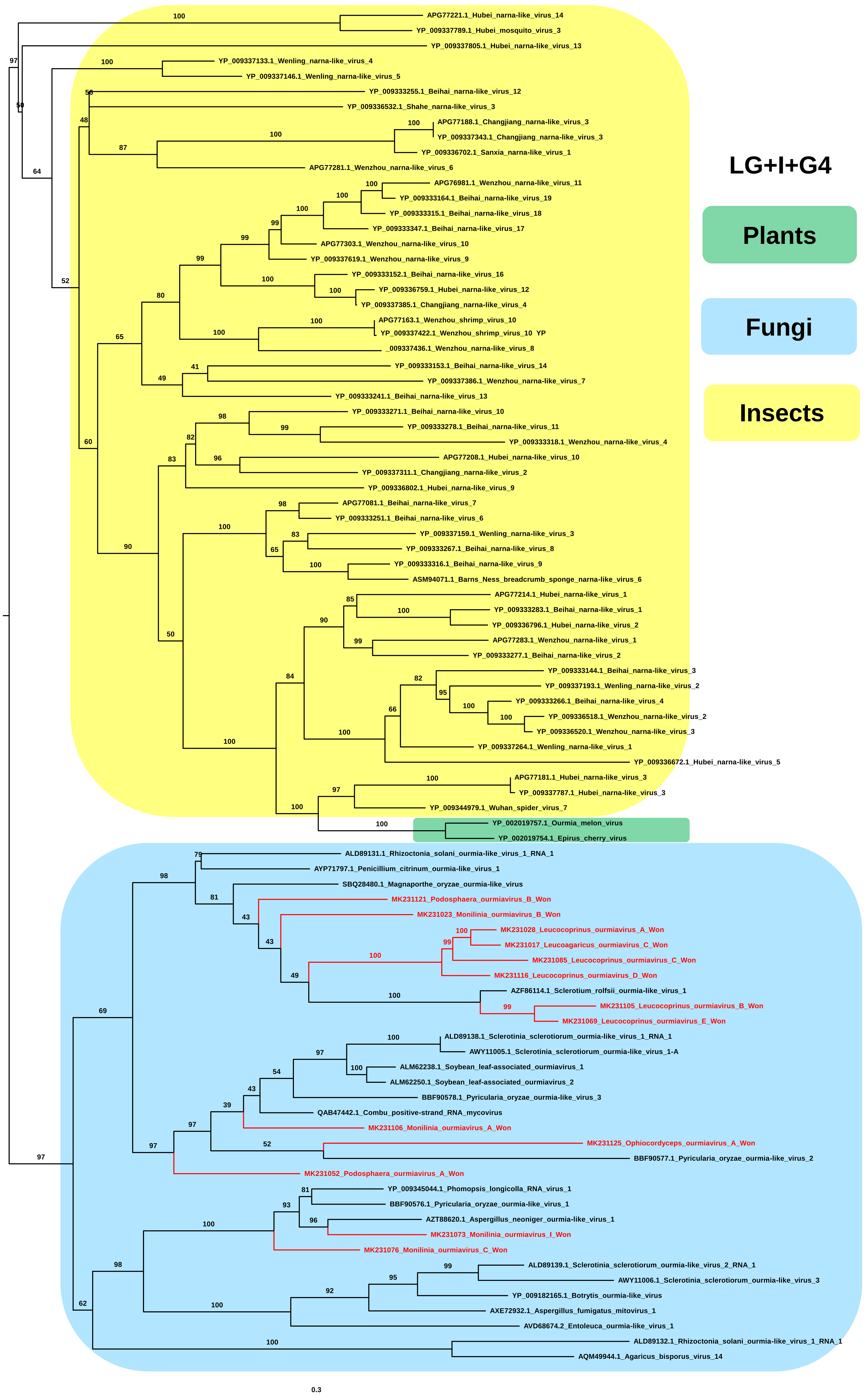


Figure S4. Phylogenetic tree of 13 viruses in family *Botourmiaviridae* and related viruses.
RdRp sequences for 13 viruses in family *Botourviridae* and related viruses were aligned with MAFFT and subjected to phylogenetic tree construction using maximum likelihood method and LG+I+G4 protein substitution model implemented in IQ-TREE. The phylogenetic tree is midpoint rooted using FigTree. Viruses in this study were indicated by red colors. Bootstrap values are indicated on branches with 1,000 replicates. Viruses derived from different hosts were indicated by different colored boxes.

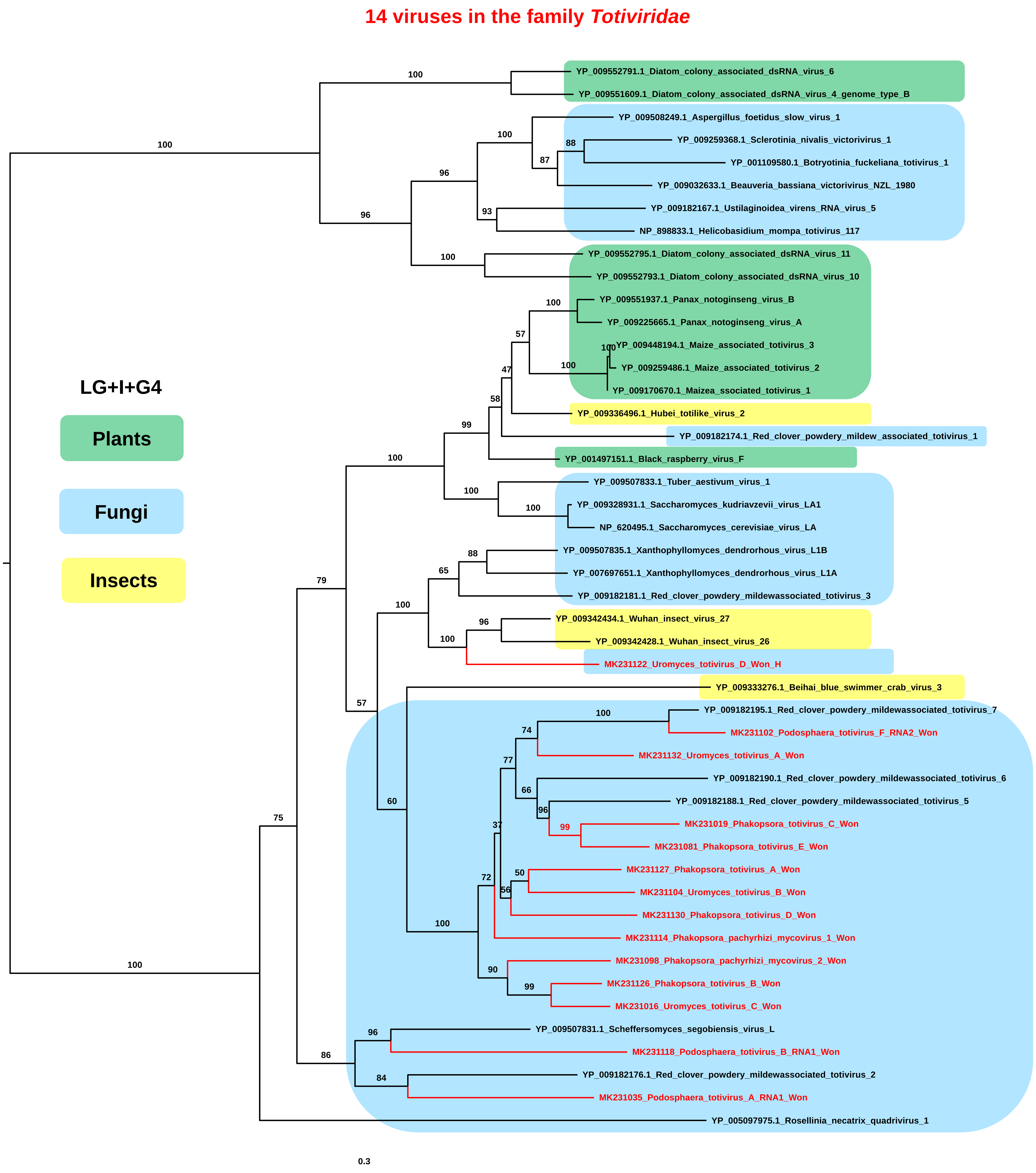


Figure S5. Phylogenetic tree of 14 viruses in family *Totiviridae* and related viruses.

RdRp sequences for 14 viruses in family *Totiviridae* and related viruses were aligned with MAFFT and subjected to phylogenetic tree construction using maximum likelihood method and LG+I+G4 protein substitution model implemented in IQ-TREE. The phylogenetic tree is midpoint rooted using FigTree. Viruses in this study were indicated by red colors. Bootstrap values are indicated on branches with 1,000 replicates. Viruses derived from different hosts were indicated by different colored boxes.

#### Nine viruses in the family *Gammaflexiviridae*

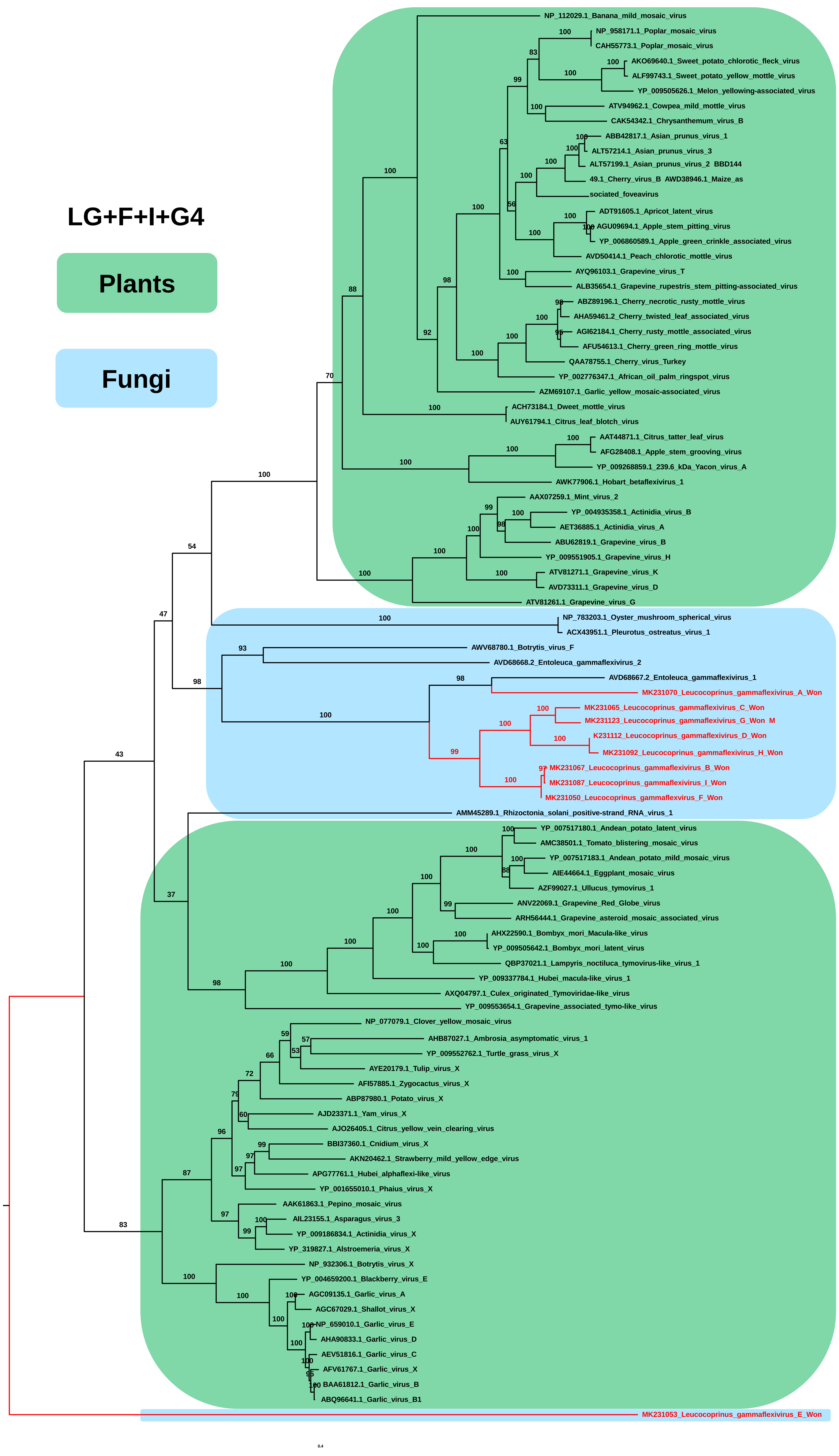


Figure S6. Phylogenetic tree of nine viruses in family *Gammaflexiviridae* and related viruses.

RdRp sequences for nine viruses in family *Gammaflexiviridae* and related viruses were aligned with MAFFT and subjected to phylogenetic tree construction using maximum likelihood method and LG+F+I+G4 protein substitution model implemented in IQ-TREE. The phylogenetic tree is midpoint rooted using FigTree. Viruses in this study were indicated by red colors. Bootstrap values are indicated on branches with 1,000 replicates. Viruses derived from different hosts were indicated by different colored boxes.

#### Seven viruses in the family *Hypoviridae*

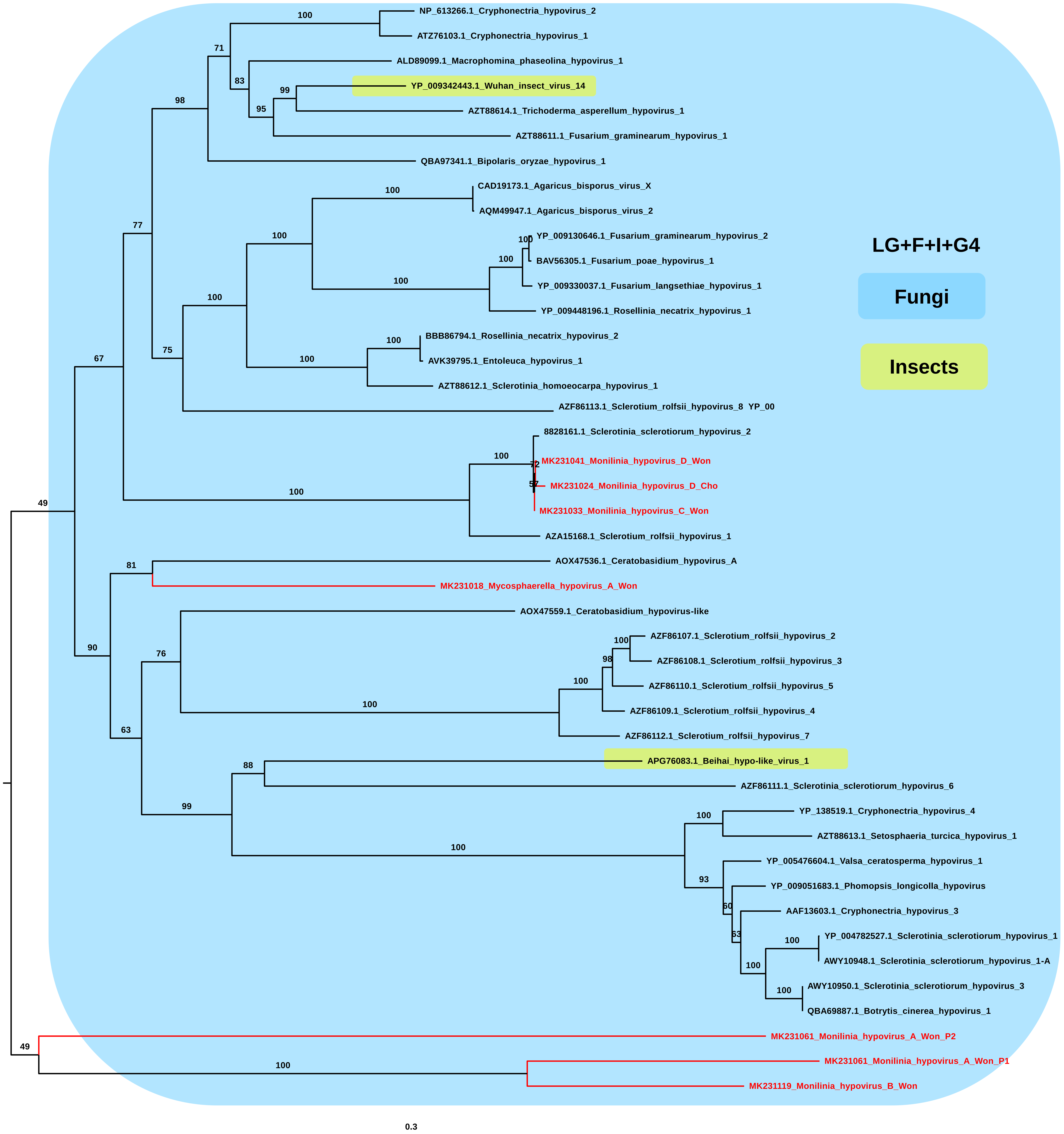


Figure S7. Phylogenetic tree of seven viruses in family *Hypoviridae* and related viruses.
RdRp sequences for nine viruses in family *Hypoviridae* and related viruses were aligned with MAFFT and subjected to phylogenetic tree construction using maximum likelihood method and LG+F+I+G4 protein substitution model implemented in IQ-TREE. The phylogenetic tree is midpoint rooted using FigTree. Viruses in this study were indicated by red colors. Bootstrap values are indicated on branches with 1,000 replicates. Viruses derived from different hosts were indicated by different colored boxes.

#### Seven viruses in the family *Partitiviridae*

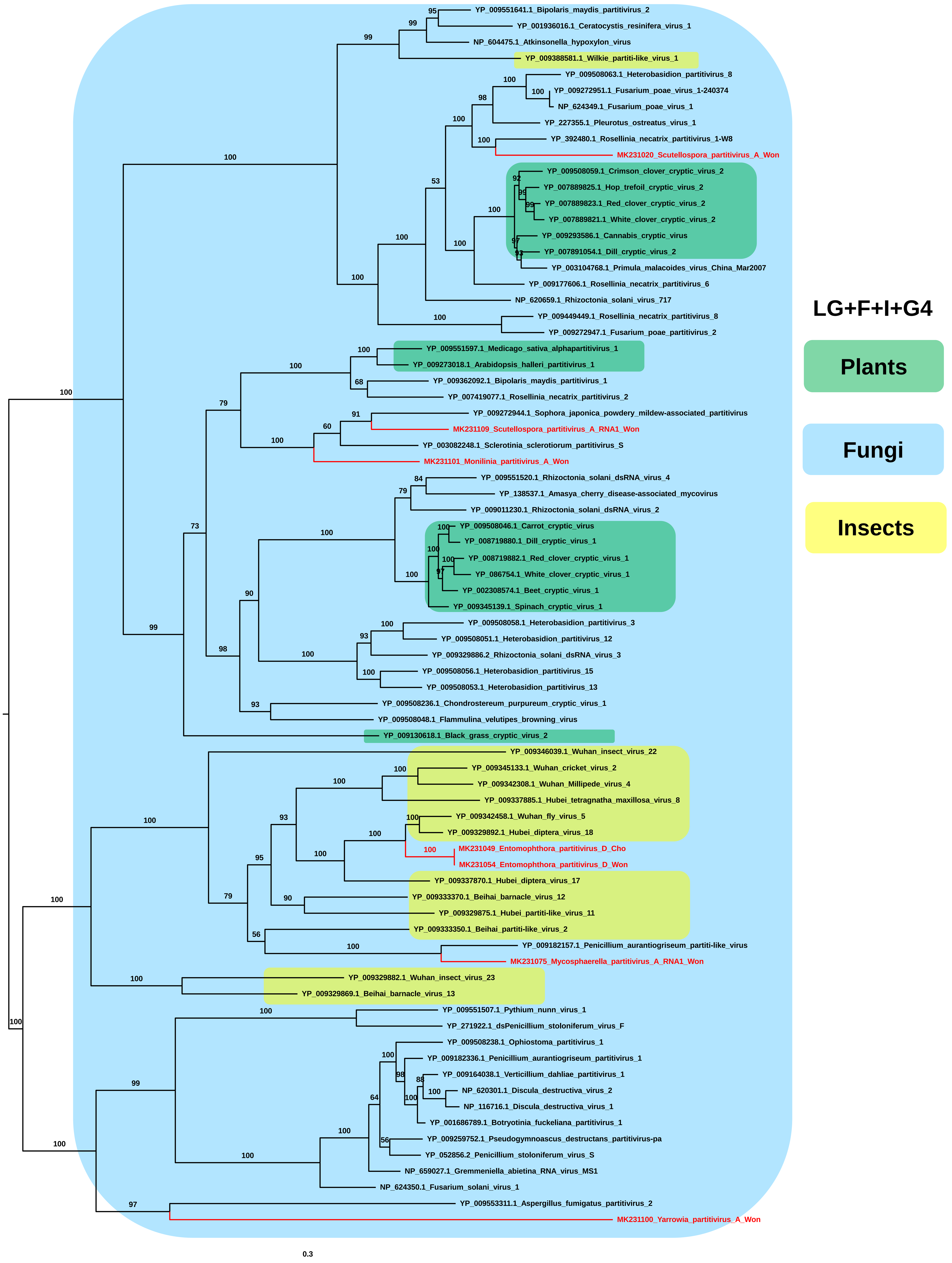


Figure S8. Phylogenetic tree of seven viruses in family *Partitiviridae* and related viruses.
RdRp sequences for seven viruses in family *Partitiviridae* and related viruses were aligned with MAFFT and subjected to phylogenetic tree construction using maximum likelihood method and LG+F+I+G4 protein substitution model implemented in IQ-TREE. The phylogenetic tree is midpoint rooted using FigTree. Viruses in this study were indicated by red colors. Bootstrap values are indicated on branches with 1,000 replicates. Viruses derived from different hosts were indicated by different colored boxes.

**Two viruses in the family *Barnaviridae***

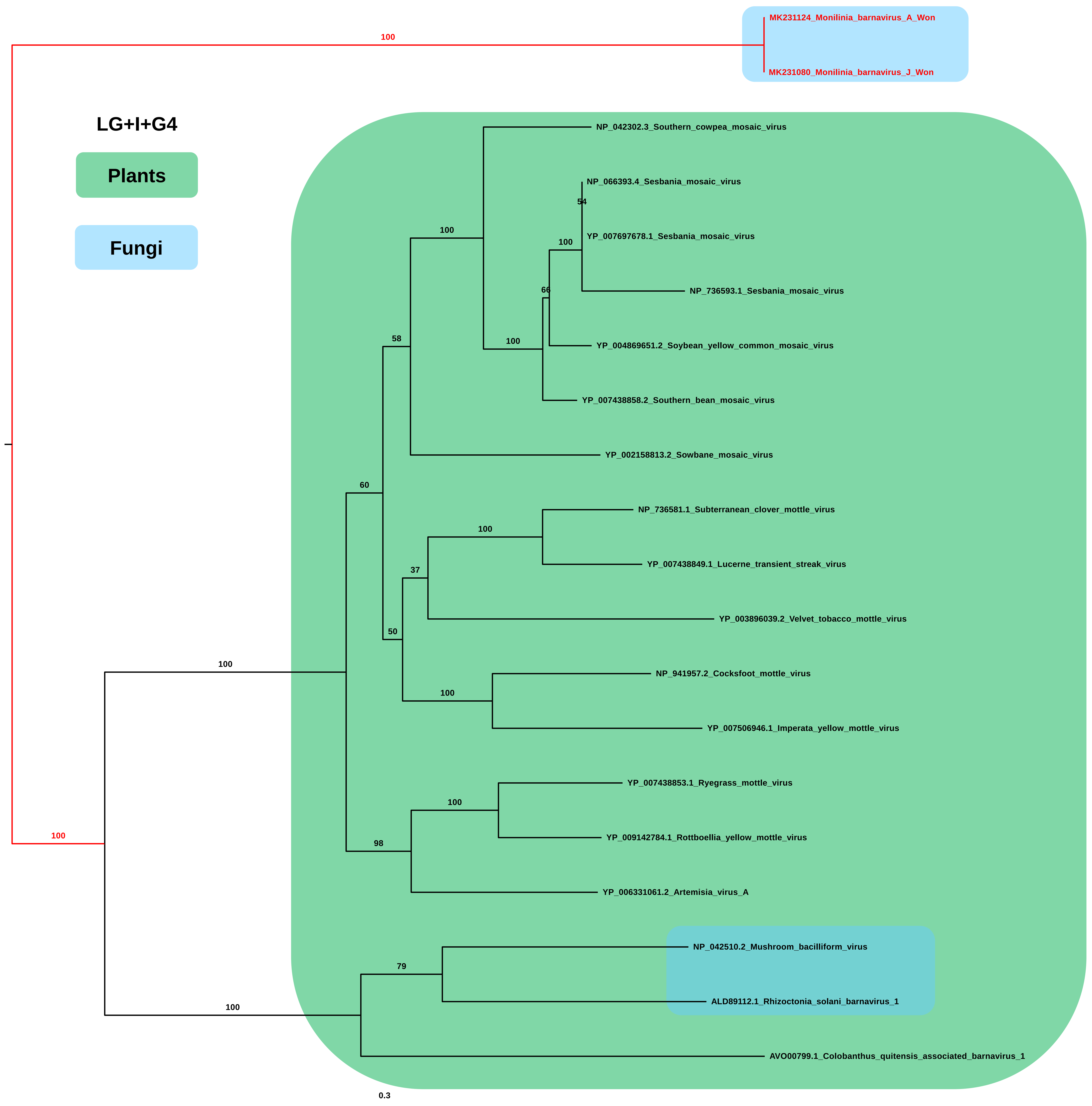


Figure S9. Phylogenetic tree of two viruses in family *Barnaviridae* and related viruses.
RdRp sequences for two viruses in family *Barnaviridae* and related viruses were aligned with MAFFT and subjected to phylogenetic tree construction using maximum likelihood method and LG+I+G4 protein substitution model implemented in IQ-TREE. The phylogenetic tree is midpoint rooted using FigTree. Viruses in this study were indicated by red colors. Bootstrap values are indicated on branches with 1,000 replicates. Viruses derived from different hosts were indicated by different colored boxes.

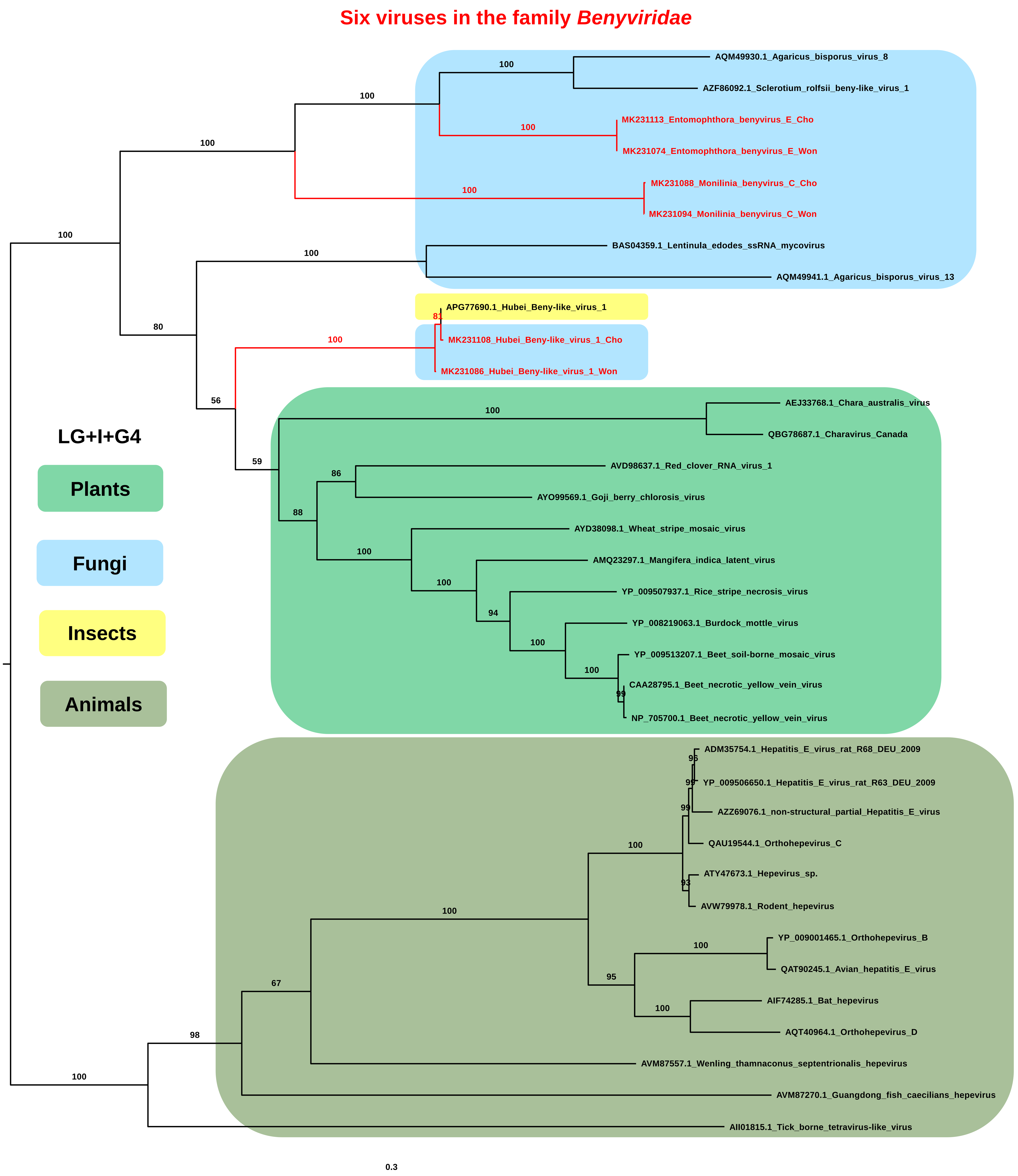


Figure S10. Phylogenetic tree of six viruses in family *Benyviridae* and related viruses.

RdRp sequences for six viruses in family *Benyviridae* and related viruses were aligned with MAFFT and subjected to phylogenetic tree construction using maximum likelihood method and LG+I+G4 protein substitution model implemented in IQ-TREE. The phylogenetic tree is midpoint rooted using FigTree. Viruses in this study were indicated by red colors. Bootstrap values are indicated on branches with 1,000 replicates. Viruses derived from different hosts were indicated by different colored boxes.

#### One virus in the family *Bromoviridae*

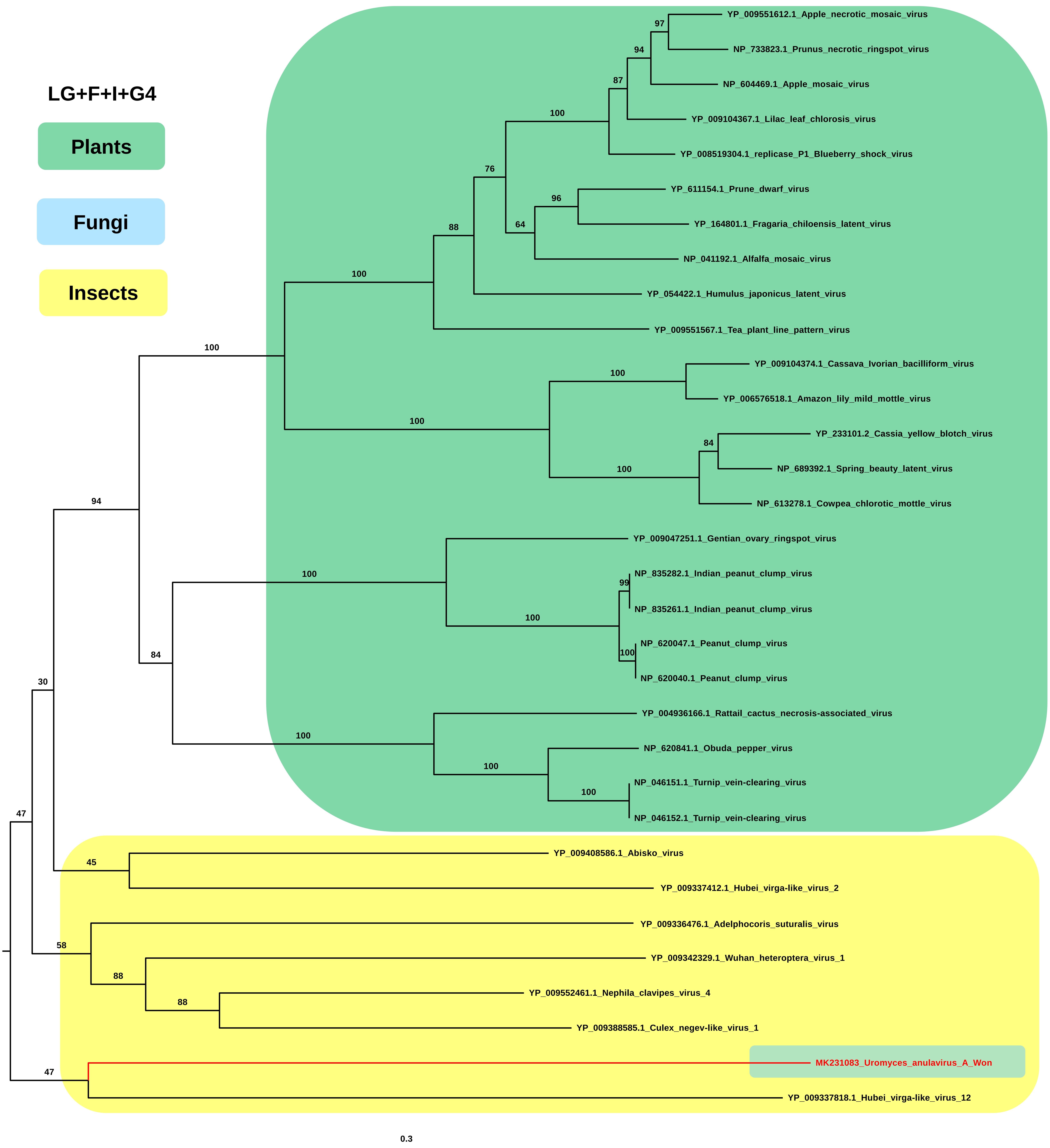


Figure S11. Phylogenetic tree of a virus in family *Bromoviridae* and related viruses.
RdRp sequences for a virus in family *Bromoviridae* and related viruses were aligned with MAFFT and subjected to phylogenetic tree construction using maximum likelihood method and LG+I+G4 protein substitution model implemented in IQ-TREE. The phylogenetic tree is midpoint rooted using FigTree. Viruses in this study were indicated by red colors. Bootstrap values are indicated on branches with 1,000 replicates. Viruses derived from different hosts were indicated by different colored boxes.

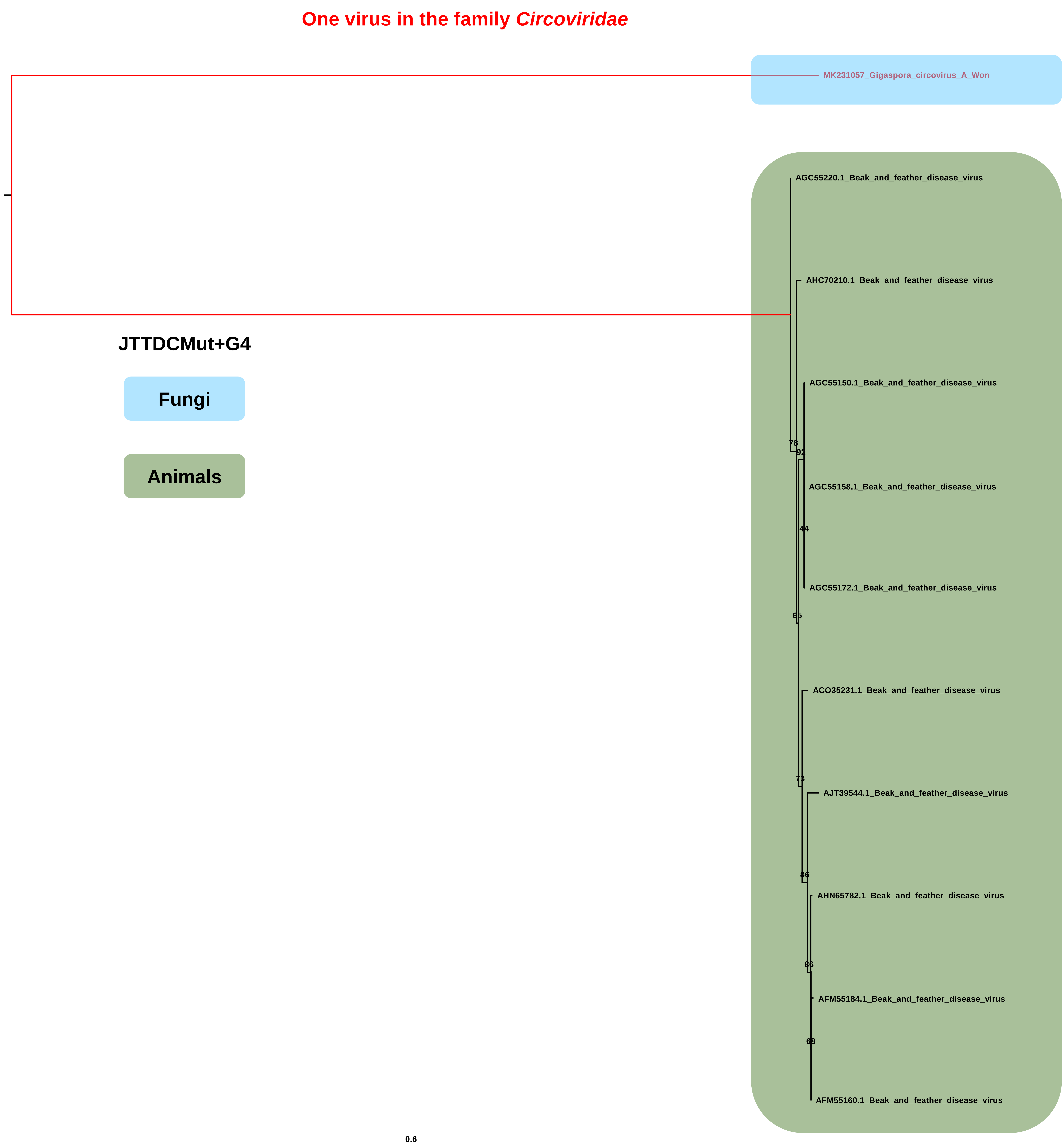


Figure S12. Phylogenetic tree of a virus in family *Circoviridae* and related viruses.
Replication initiator protein sequences for a virus in family *Circoviridae* and related viruses were aligned with MAFFT and subjected to phylogenetic tree construction using maximum likelihood method and the JTTDCMut+G4 protein substitution model implemented in IQ-TREE. The phylogenetic tree is midpoint rooted using FigTree. Viruses in this study were indicated by red colors. Bootstrap values are indicated on branches with 1,000 replicates. Viruses derived from different hosts were indicated by different colored boxes.

#### Two viruses in the family *Fusaviridae*

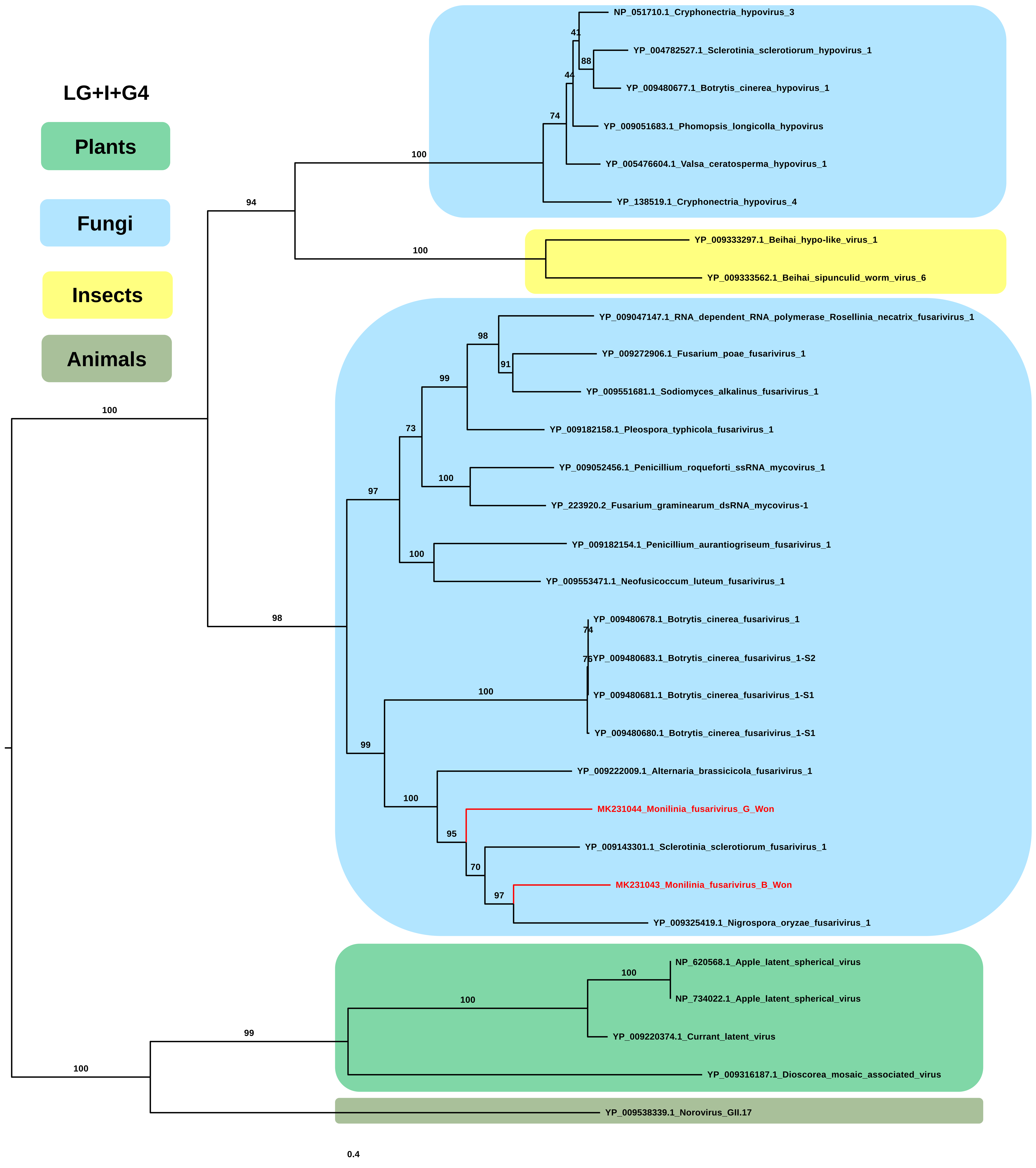


Figure S13. Phylogenetic tree of two viruses in family *Fusariviridae* and related viruses.
RdRp sequences for two viruses in family *Fusariviridae* and related viruses were aligned with MAFFT and subjected to phylogenetic tree construction using maximum likelihood method and LG+I+G4 protein substitution model implemented in IQ-TREE. The phylogenetic tree is midpoint rooted using FigTree. Viruses in this study were indicated by red colors. Bootstrap values are indicated on branches with 1,000 replicates. Viruses derived from different hosts were indicated by different colored boxes.

#### One virus in the family *Phasmaviridae*

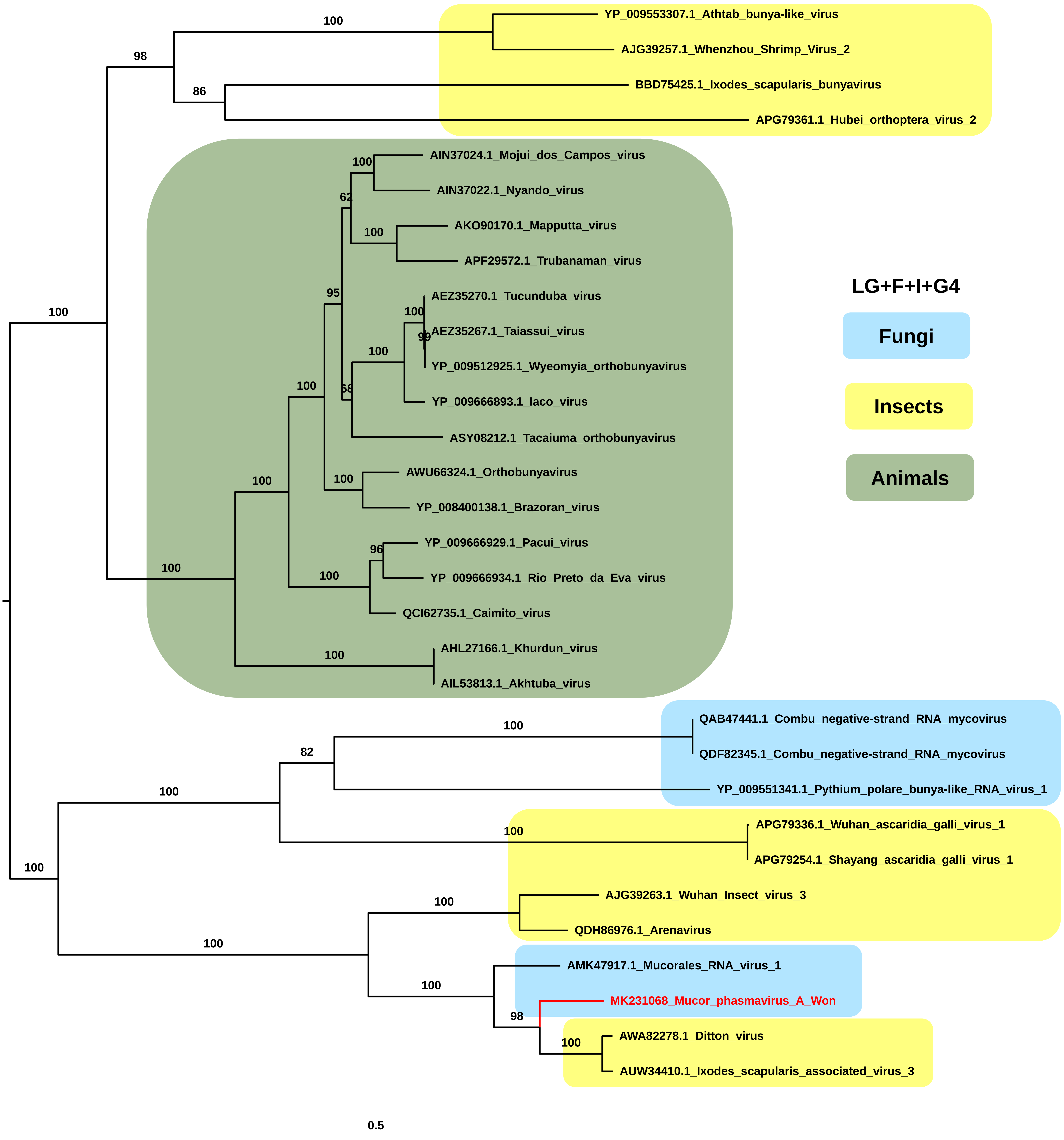


Figure S14. Phylogenetic tree of a virus in the family *Phasmaviridae* and related viruses.
RdRp sequences for a virus in the family *Phasmaviridae* and related viruses were aligned with MAFFT and subjected to phylogenetic tree construction using maximum likelihood method and LG+F+I+G4 protein substitution model implemented in IQ-TREE. The phylogenetic tree is midpoint rooted using FigTree. Viruses in this study were indicated by red colors. Bootstrap values are indicated on branches with 1,000 replicates. Viruses derived from different hosts were indicated by different colored boxes.

#### Two viruses in the order *Picornavirales*

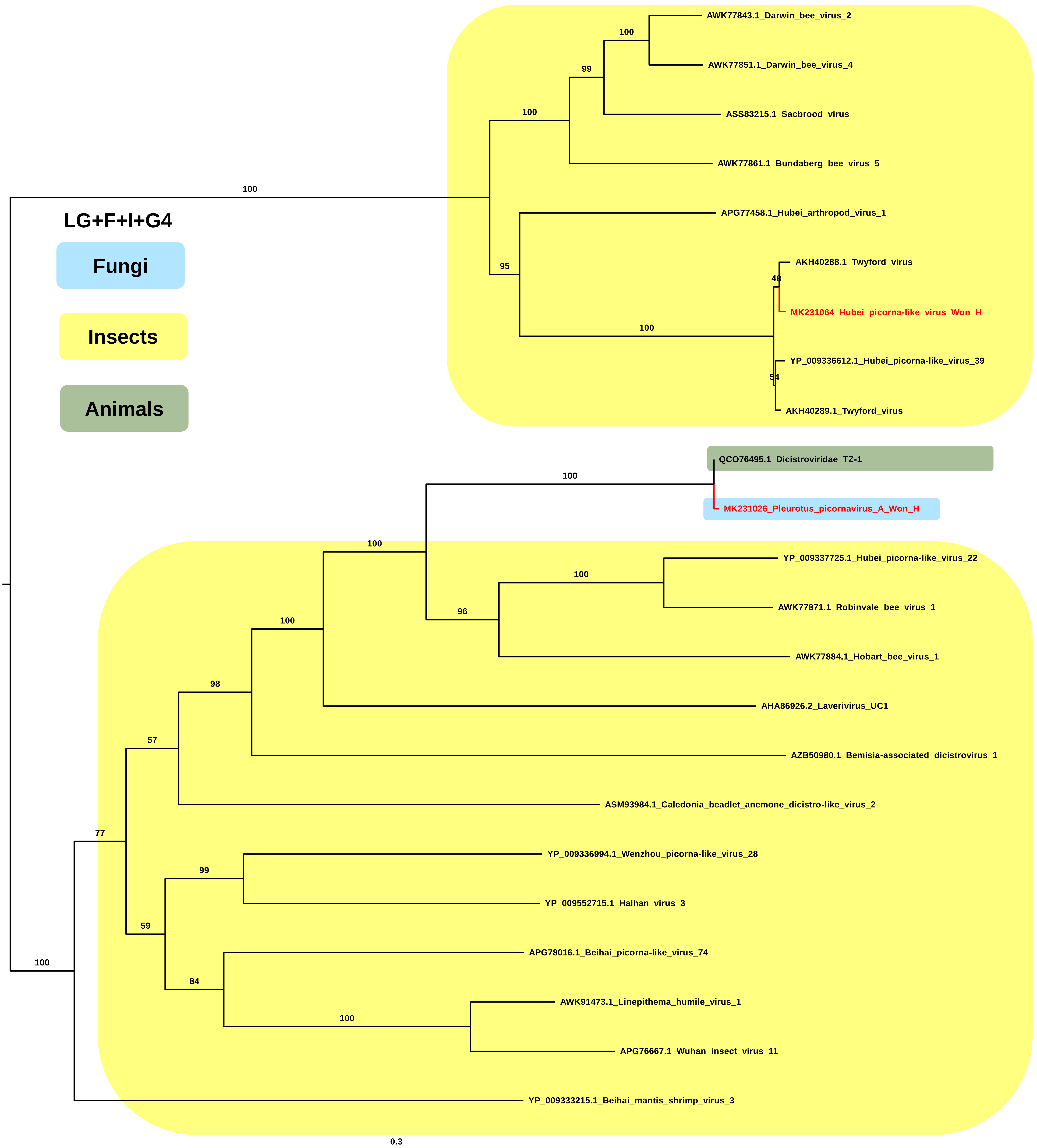


Figure S15. Phylogenetic tree of two viruses in the order *Picornavirales* and related viruses. RdRp sequences for two viruses in the order *Picornavirales* and related viruses were aligned with MAFFT and subjected to phylogenetic tree construction using maximum likelihood method and LG+F+I+G4 protein substitution model implemented in IQ-TREE. The phylogenetic tree is midpoint rooted using FigTree. Viruses in this study were indicated by red colors. Bootstrap values are indicated on branches with 1,000 replicates. Viruses derived from different hosts were indicated by different colored boxes.

One virus in the family *Potyviridae*

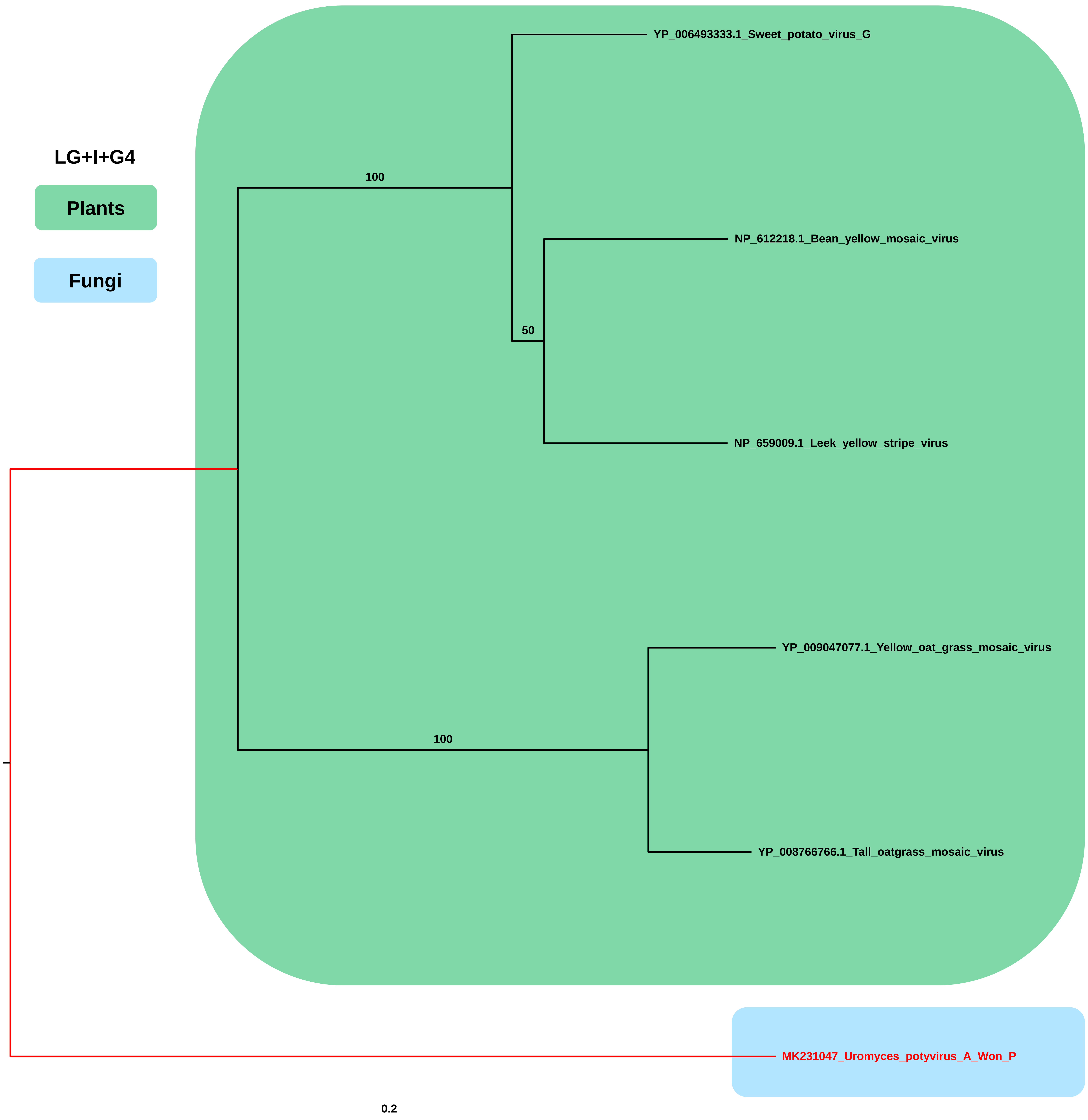


Figure S16. Phylogenetic tree of a virus in the family *Potyviridae* and related viruses.
RdRp sequences for a virus in the family *Potyviridae* and related viruses were aligned with MAFFT and subjected to phylogenetic tree construction using maximum likelihood method and LG+I+G4 protein substitution model implemented in IQ-TREE. The phylogenetic tree is midpoint rooted using FigTree. Viruses in this study were indicated by red colors. Bootstrap values are indicated on branches with 1,000 replicates. Viruses derived from different hosts were indicated by different colored boxes.

#### One virus in the family *Secoviridae*

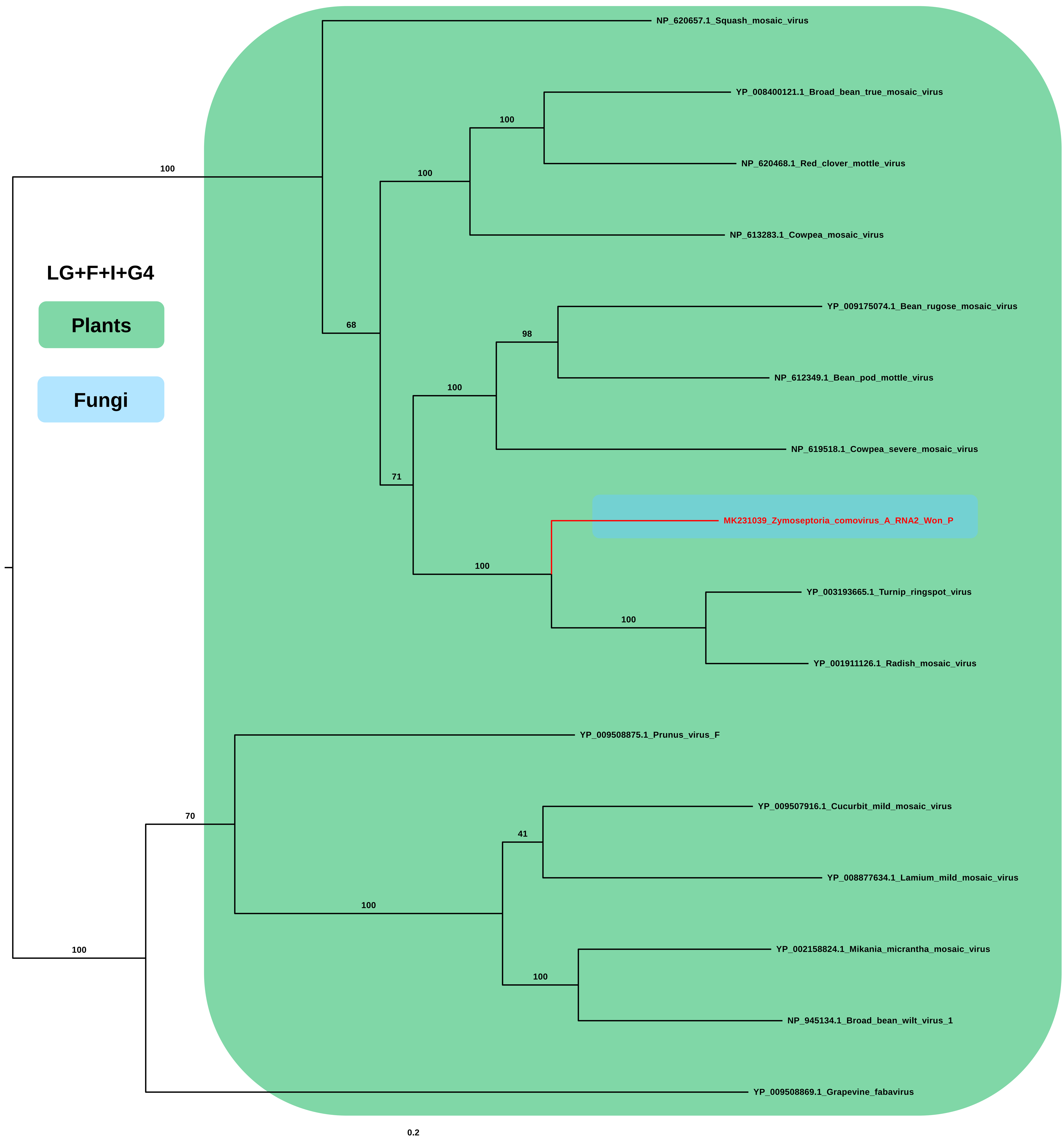


Figure S17. Phylogenetic tree of a virus in the family *Secoviridae* and related viruses.
RdRp sequences for a virus in the family *Secoviridae* and related viruses were aligned with MAFFT and subjected to phylogenetic tree construction using maximum likelihood method and LG+F+I+G4 protein substitution model implemented in IQ-TREE. The phylogenetic tree is midpoint rooted using FigTree. Viruses in this study were indicated by red colors. Bootstrap values are indicated on branches with 1,000 replicates. Viruses derived from different hosts were indicated by different colored boxes.

### One virus in the family *Rhabdoviridae*

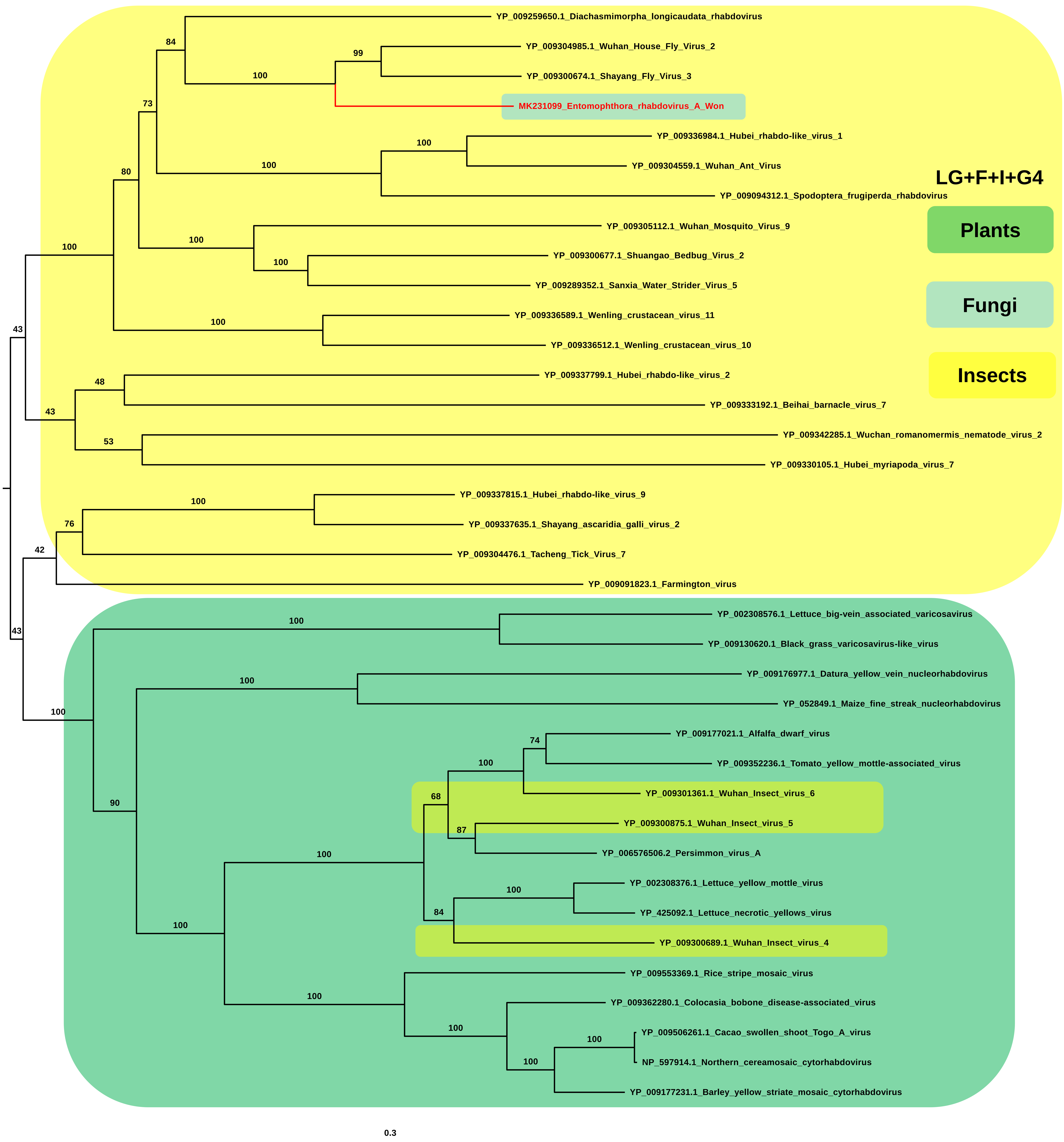


Figure S18. Phylogenetic tree of a virus in the family *Rhabdoviridae* and related viruses.
RdRp sequences for a virus in the family *Rhabdoviridae* and related viruses were aligned with MAFFT and subjected to phylogenetic tree construction using maximum likelihood method and LG+F+I+G4 protein substitution model implemented in IQ-TREE. The phylogenetic tree is midpoint rooted using FigTree. Viruses in this study were indicated by red colors. Bootstrap values are indicated on branches with 1,000 replicates. Viruses derived from different hosts were indicated by different colored boxes.

#### Three viruses in the family *Tymoviridae*

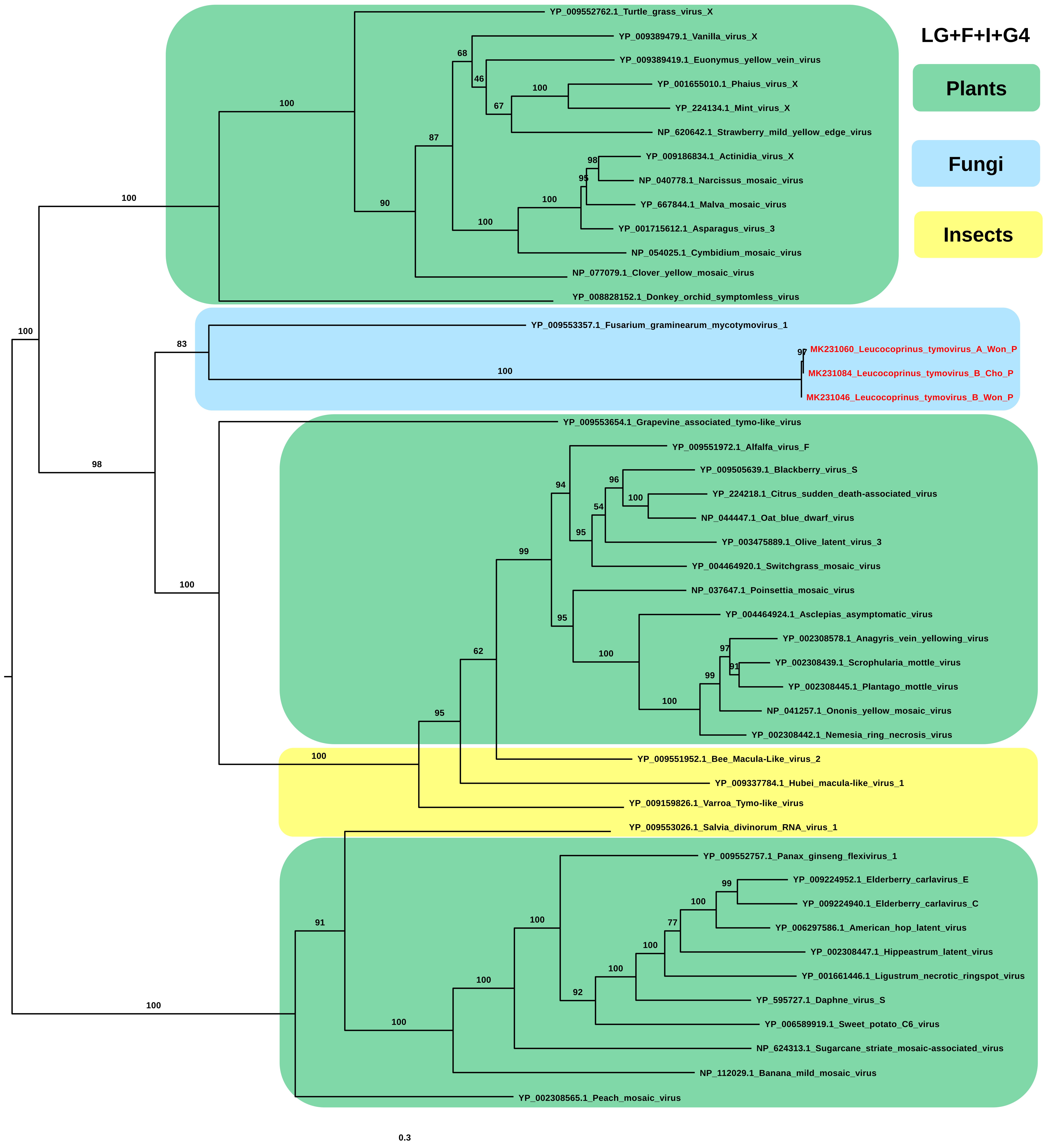


Figure S19. Phylogenetic tree of three viruses in the family *Tymoviridae* and related viruses.
RdRp sequences for three viruses in the family *Tymoviridae* and related viruses were aligned with MAFFT and subjected to phylogenetic tree construction using maximum likelihood method and LG+F+I+G4 protein substitution model implemented in IQ-TREE. The phylogenetic tree is midpoint rooted using FigTree. Viruses in this study were indicated by red colors. Bootstrap values are indicated on branches with 1,000 replicates. Viruses derived from different hosts were indicated by different colored boxes.

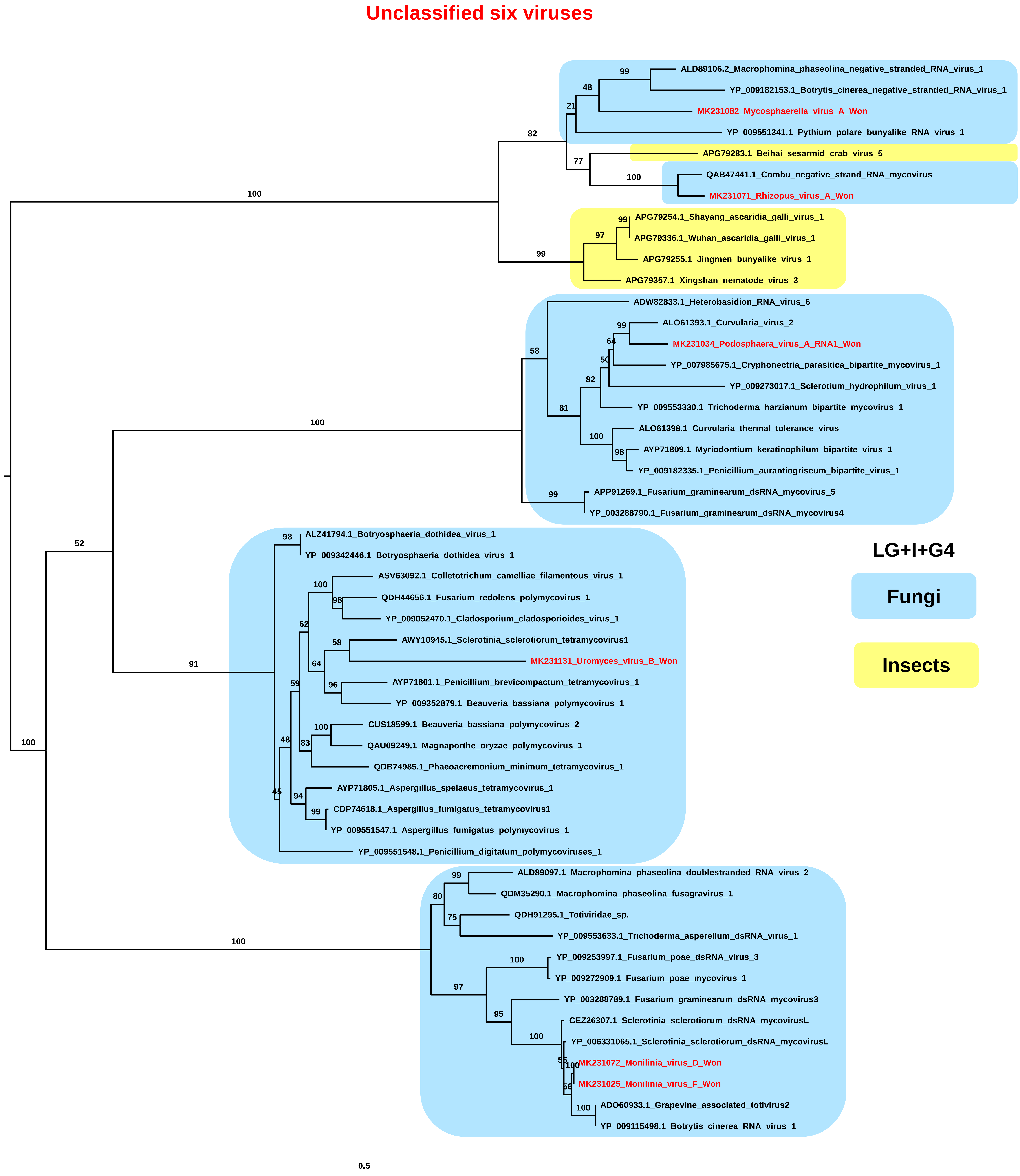


Figure S20. Phylogenetic tree of six unclassified viruses and related viruses.
RdRp sequences for six unclassified viruses and related viruses were aligned with MAFFT and subjected to phylogenetic tree construction using maximum likelihood method and LG+I+G4 protein substitution model implemented in IQ-TREE. The phylogenetic tree is midpoint rooted using FigTree. Viruses in this study were indicated by red colors. Bootstrap values are indicated on branches with 1,000 replicates. Viruses derived from different hosts were indicated by different colored boxes.
